# Phenotyping Polarization Dynamics Of Immune Cells Using A Lipid Droplet - Cell Pairing Microfluidic Platform

**DOI:** 10.1101/2021.12.22.473360

**Authors:** Léa Pinon, Nicolas Ruyssen, Judith Pineau, Olivier Mesdjian, Damien Cuvelier, Anna Chipont, Rachele Allena, Coralie L. Guerin, Sophie Asnacios, Atef Asnacios, Paolo Pierobon, Jacques Fattaccioli

## Abstract

The immune synapse is the tight contact zone between a lymphocyte and a cell presenting its cognate antigen. This structure serves as a signaling platform and entails a polarization of intra-cellular components, necessary to the immunological function of the cell. While the surface properties of the presenting cell are known to control the formation of the synapse, their impact on polarization has not yet been studied.

Using functional lipid droplets as tunable artificial presenting cells combined with a microfluidic pairing device, we simultaneously observe synchronized synapses and dynamically quantify polarization patterns of individual B cells. By assessing how ligand concentration, surface fluidity and substrate rigidity impact lysosome polarization, we show that its onset and kinetics depend on the local antigen concentration at the synapse and on substrate rigidity. Our experimental system enables a fine phenotyping of monoclonal cell populations based on their synaptic readout.

## Introduction

Direct contact is an important channel of communication for cells in multicellular organisms. This is true for cells in tissues as well as for cells that mostly live as independent entities like immune cells (1). Their activation, their immune function and ultimately their fate depend on signal exchanges with other cells through an organized structure called immune synapse. In both B and T lymphocytes, the formation of the immune synapse is associated with a global rearrangement of the cytoskeleton and the establishment of a polarity axis (2–4). B lymphocytes, the cells responsible for antibody production, encounter antigens in the subcapsular sinus of the lymph node, in soluble form or grafted on other cell surface such as macrophages or follicular dendritic cells. They recognize the antigen through their specific B cell receptor (BCR), internalize, process and further present it to a cognate T cell. Antigen recognition entails membrane and intracellular reorganizations leading to the formation of the immune synapse (5). Engagement of the BCR leads to the clusterization of signaling complexes in a mechanosensitive way: signaling and size of the complexes depend on the rigidity and topography of the substrate (6–8). The synapse, in its final form, displays a stereotypical concentric shape with antigens/BCRs accumulated in a central cluster, surrounded by adhesion molecules and an actin ring at the periphery (9—11). A similar geometry is mirrored by cytoplasmic molecules close to the membrane (12). At the same time, the microtubule network re-orients the traffic towards the synapse (polarization of the centrosome) to secrete lysosomal proteases and degrade the antigen in the synaptic cleft, in a process named enzymatic extraction (13). It has been shown that, depending on the deformability of the substrate, the antigen can be internalized also by mechanical pulling (14—17). When mechanical extraction fails, such as on non-deformable substrates, cells trigger the enzymatic extraction pathways described above (15). Interestingly, the mutual exclusivity of mechanical and enzymatic extractions suggests that the polarization mechanism is also sensitive to the mechanical properties of the substrate. Ultimately, the B cell immune synapse results in signal transduction, cell differentiation and production of high-affinity antibodies (2). It has been also proposed that by polarizing, B cells can divide asymmetrically to give rise to B cells that present more efficiently the antigen to T cells. Polarity, therefore, has consequences on B cell fate (18, 19).

All these experimental results point to a crucial role of physico-chemical properties of the antigen presenting surface in B cell activation. Different systems have been used to address this mechanism. For instance, clusters formation has been revealed on fluid interfaces allowing antigen mobility (lipid bilayers (9)); mechanosensitivity has been shown using deformable substrates such as soft gels (6, 7); antigen mechanical extraction has been uncovered on plasma membrane sheets (14), and quantified by calibrated DNA force sensors (15, 20). Despite the amount of information gathered in these systems, the variety of assays hinders the comparison between experiments performed on different materials and makes it impossible to evaluate the impact of independent properties on the synapse formation. This prompts us to introduce a new model to stimulate B cells while independently controlling physical (rigidity, fluidity, size) and chemical (functionalization) properties: lipid droplets.

Emulsions are colloidal liquid-liquid metastable suspensions stabilized by a surfactant monolayer, that have already shown their biocompatibility and their interest as probes when functionalized with proteins of interests in biophysical (21), developmental (22), and immunological contexts such as phagocytosis (23—25) or T cell synapse studies (26). By varying the bulk and surface composition, it is possible to tune the surface tension, hence the mechanical rigidity, independently from the ligand surface concentration, thus making lipid droplets a relevant antigen-presenting cell (APC) surrogate to stimulate B cells with the highest control on the physico-chemical properties of the cognate surface.

In this work, we introduce new droplet formulations and functionalization to access different physical and chemical properties that we finely characterize. Finally, we validate our methods n addressing how cells polarize depending on the properties of those antigen-presenting substrates. This question has been neglected in the past, partially because of the heterogeneity of experimental models, partially because of the lack of reproducible way to study the global cell rearrangement following immune synapse formation over time. Therefore, for a proper quantitative study, we engineered antigen-functionalized lipid droplets with a pertinent set of physico-chemical properties and presented them to B cells in a controlled microfluidic pairing device that minimizes the stress to mimic the flow conditions of the lymph node. This allowed us to simultaneously observe multiple synchronized synapses with a high spatio-temporal resolution and finally provide a phenotyping map of variability of B cell polarity both in terms of polarization onset and kinetics.

## Results

### Microfluidic cell-cell pairing platform for the study of B cell synapses

To characterize the influence of rigidity and antigen concentration on B cell polarity, we observe by epifluorescence microscopy the polarization of murine B cells activated by antigen-coated fluid microparticles having different mechanical and interfacial properties.

Because B cells are non-adherent cells, to control in space and time B cell synapse, we used a microfluidic chip that forces the encounter between a single B cell and an activating APC surrogate (**Fig. 1A**). The microfluidic chip (**Fig. 1B**), inspired by (27), consists of staggered arrays of double-layered U-shaped traps where the two objects (ideally of the same size) are sequentially immobilized (**Supplementary Note 1**) and imaged. The double layer structures result in fluid streamlines not being deviated from the weir structures when a first object is trapped, thus allowing the easy capture of a second object (27, 28). The trapping array was engineered to apply the least perturbative shear stress as possible on the cells, mimicking the *in vivo*-like shear stress estimated in the subcapsular sinus lumen of lymphoid tissues (29). Finite element simulations (**Supplementary Note 1**) indicate that, at the inlet speed used in our experiments, the maximal wall shear stresses on wall (**Fig. 1C**) and trapped cell (**Fig. 1D**) are below 1 Pa (value compatible with the shear stress estimated in the lymph node (29)). Overall, the microfluidic chip we designed minimizes the stress to mimic conditions that a cell experiences in lymph nodes.

**Fig. 1.**
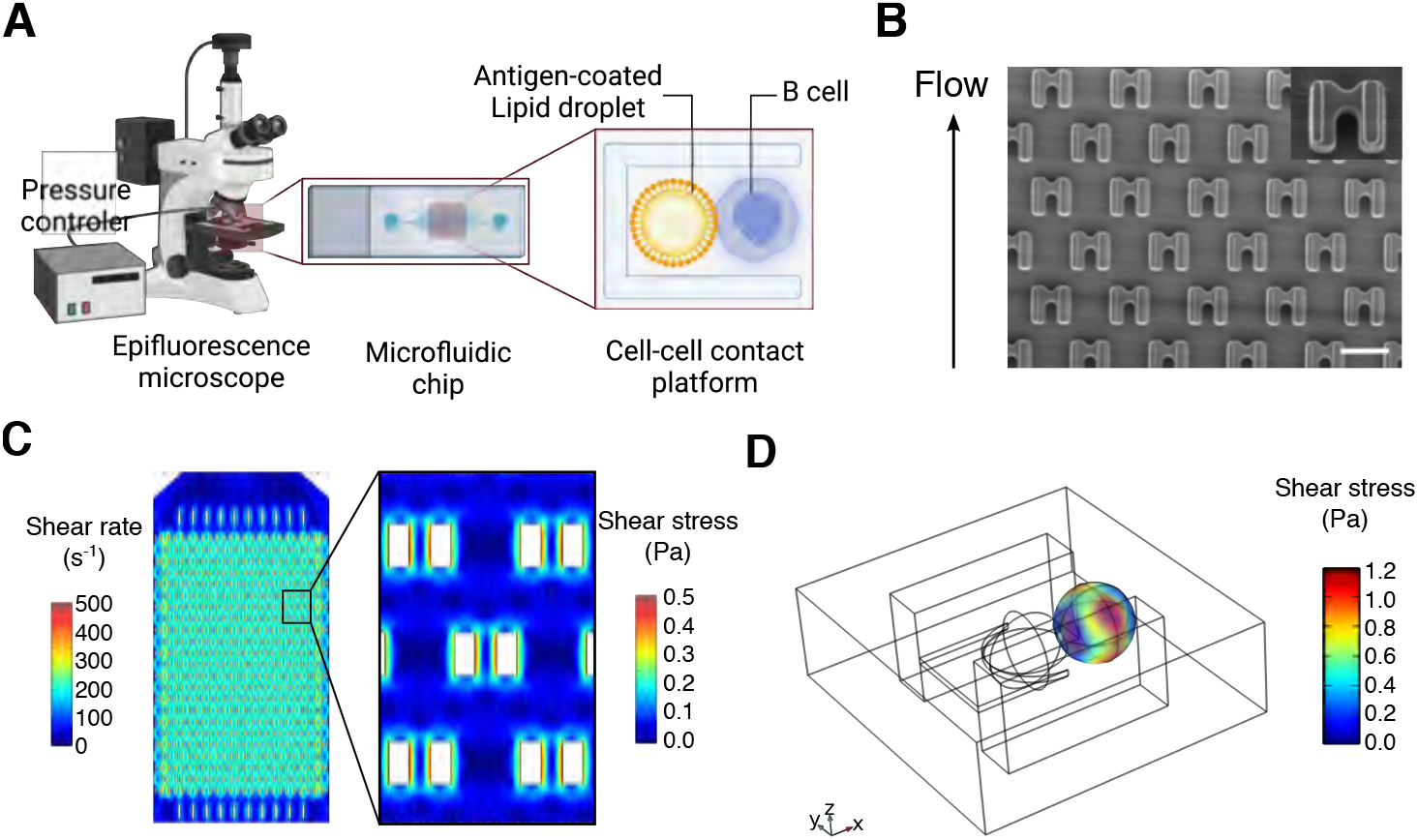
Microfluidic cell-cell pairing platform for the study of B cell synapses. (A) Schematic view of the experimental setup. The microfluidic trap-based chip is imaged with an epifluorescence video-microscope, and is connected to a pressure controller inducing a low flow for droplets - then cells - circulation. (B) SEM images of the microfluidic chamber containing 288 two-layered traps. Scale bar: 30 *μm*. (C) 2D FEM simulation of fluid shear rate (left) and shear stress (right) passing through the pillars along the whole chip, for an inlet pressure of 1000 Pa corresponding to a maximal fluid velocity of 1.6 mm/s. The shear rate is constant along the chamber. The maximal wall shear rate is about 514 s^-1^ that corresponds to a maximal wall shear stress about 0.514 Pa. (D) 3D simulation of the shear stress that a trapped cell experiences in the microfluidic device when it is immobilized with a droplet (wireframe representation). For a maximal inlet pressure of 1000 Pa - maximal fluid velocity of 1.6 mm/s in the chamber - the maximal shear stress is about 1.0 Pa.

### Droplets mimic APC both in antigen concentrations and mechanical properties

Our study has been conducted with B lymphoma murine cell line IIA1.6, which is described to be a homogeneous population of non-activated mature B cells. These cells constitute an ideal model for functional studies of antigen presentation, since they lack Fc*γ*RIIB1 receptors on their surface (30, 31) and hence can be conveniently activated through specific antibodies against the BCR (F(ab’)_2_). We stimulated these cells with lipid droplets functionalized with such antibodies (which will be referred to as antigens, Ag), linked to droplet surface with biotinylated phospholipids (DPSE-PEG_2000_-Biotin)/streptavidin complex **Fig. 2A**. As negative control, droplets were coated with biotinylated BSA which do not engage the BCR, in lieu of F(ab’)_2_ (17). In both cases, fluorescent streptavidin linker allows to observe the functionalization (**Fig. 2B**).

**Fig. 2.**
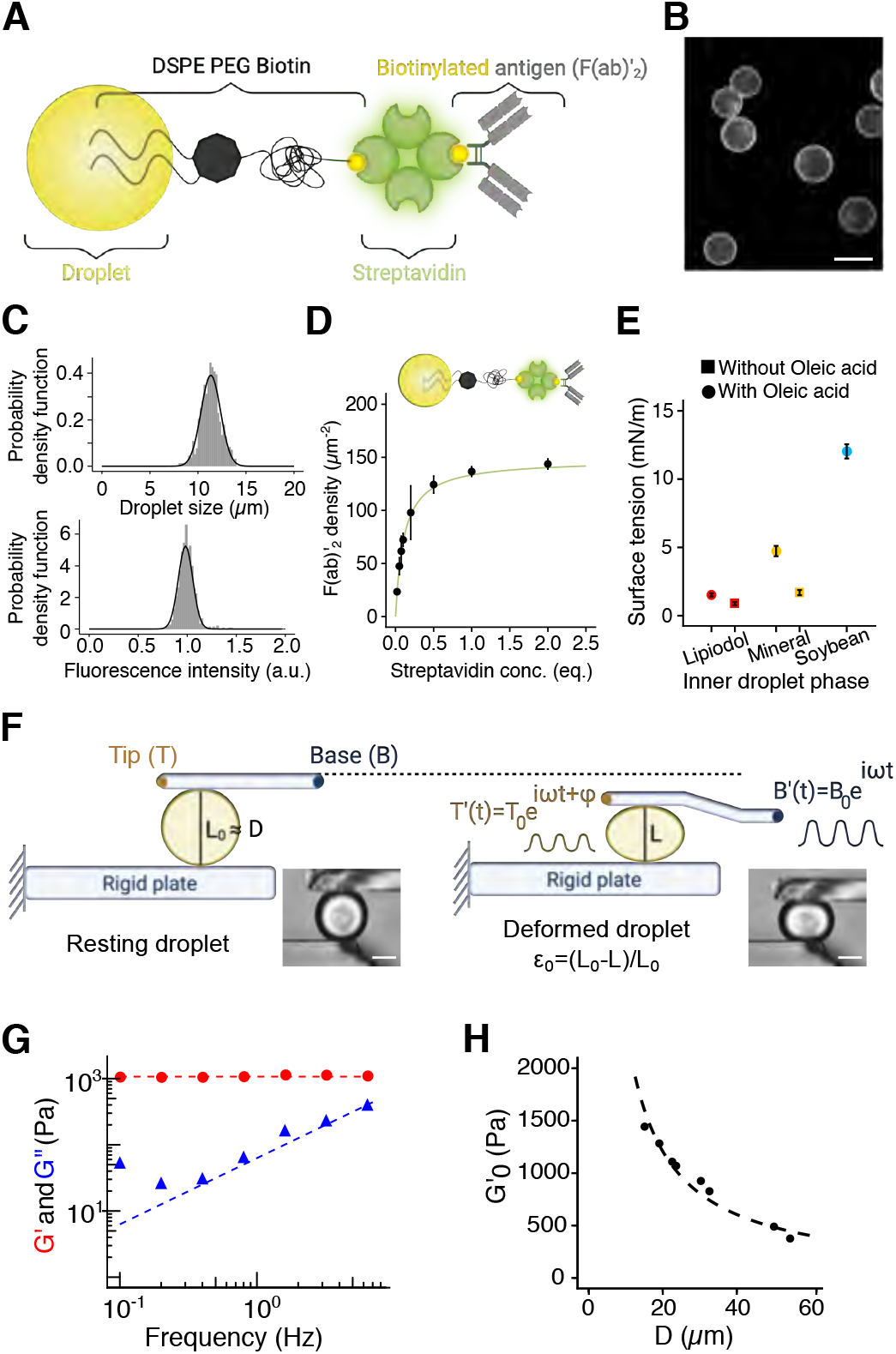
Droplets mimic APC both in antigen concentrations and mechanical properties. (A) Schematic of the droplet coating complex with DSPE-PEG_2000_-Biotin / Streptavidin / F(ab’)_2_. (B) Representative epifluorescence image of coated droplet. Scale bar: 7 *μ*m. (C) Size (mean ± std= 11.0 ± 0.7 *μ*m) and fluorescence (mean ± std = 1 ± 0.075) histograms of N = 700 droplets. (D) Titration curve of F(ab’)_2_ fragments depending on the streptavidin concentration on droplets. The fit follows to the Langmuir’s isotherm with a plateau at 147 F(ab’)_2_/μm^-2^. These data are obtained from previous fluorescence intensity assessments of both streptavidin and F(ab’)_2_ on 11 *μ*m-large droplets, initially functionalized with 100 equivalents of DSPE-PEG_2000_-biotin. (E) Pendant drop measurements of surface tension for three different oil mixtures - lipiodol (red), mineral (yellow) and soybean (blue) oils, enriched (dots) or not (squares) with oleix acid. (F) Schematic of the microplate experiment. A cell or a droplet is trapped between two glass microplates, one being immobile (bottom) the other flexible (top) with a base position oscillating as B(*t*) = *B*_0_e^*iωt*^. We measure the oscillating displacement of the tip of the flexible plate as T(*t*) =T_0_*e*^(*iωt*+*ϕ*)^ that relates to the immobilized cell or droplet visco-elastic properties. T is determined for a frequency range between 0.1 Hz and 6.4 Hz (droplet) or 1.6 Hz (cells). Representative brightfield images of a resting (left) and a deformed (right) droplet. Scale bars: 30 *μ*m. (G) Elastic G’ and viscous G″ moduli of a single droplet (diameter: 19.1 *μ*m) as a function of the probing frequency. In this frequency range, *G*’ is constant, whereas G″ linearly depends on frequency. (H) Elastic modulus of droplets plotted as a function of the resting droplet diameter, for a constant initial deformation *ϵ*_0_ = 0.2. The fitting equation is written as *G*’ = 4*γ/ϵL*_0_ and leads to a surface tension *γ* = 1.21 ± 0.04 mN.m^-1^.

Lipid droplets were fabricated by shearing an oil phase in an aqueous buffer containing surface active agents (surfactants) to improve suspension stability over time (32). To avoid a possible bias in the analysis of the synapse formation and cell polarization, that could exist if cells were put into contact with particles of different diameters (hence curvature), we decided to work with monodisperse size distributions with an average diameter of 11 *μm*, comparable to the one of a B cell (**Fig. 2B-C**).

After droplet emulsification, we coated the droplet surface by adsorbing phospholipids onto the surface with the help of a polar co-solvent, as done in (33). Then, fluorescent streptavidins, used as linker, was added to attach on the one side to the biotinylated phospholids, and bind to the other side to biotinylated F(ab’)_2_, added in the last step. This protocol ensures a finely-controlled and homogeneous lipid coating. Ultimately, protein functionalization of the droplets surface (streptavidin) is performed, as shown in **Fig. 2B** and **C**. To quantify the surface concentration in F(ab’)_2_, we used specific and fluorescent secondary antibodies at saturating concentrations and correlated streptavidin fluorescence intensity to assess the absolute F(ab’)_2_ concentration over droplet surface (**Supplementary Note 2**). **Fig. 2D** shows that the F(ab’)_2_ adsorption is well described by a Langmuir isotherm, making straightforward the quantification of their surface density (**Supplementary Note 2**). F(ab’)_2_ surface density, expressed as a number of proteins per unit of surface area, has a maximal value of 150 antigens/*μ*m^2^ at saturating conditions. This value has been previously shown to be sufficient to trigger B cell activation (10).

Antigen-presenting cells stiffness has been reported to be a relevant mechanical property for immune cell activation (34, 35) and in particular for B cell functions (6, 15). The stiffness of a material is related to its ability to resist reversible deformations when submitted to stress, i.e. to its apparent elastic properties. For oil droplets, the origin of elastic-like resistance to deformation is expected to be the excess pressure inside the droplet, i.e. the Laplace pressure ΔP written as ΔP = 2*γ*/*R*, where R is the droplet radius. The stiffness of a droplet can, therefore, be modulated by changing the surface tension *γ* between the oil and the surrounding liquid medium. We measured the surface tension of diverse droplet formulations by the pendant drop technique and micropipette (36, 37) (**Fig. S1**). We found that the interfacial tension of the oil/water interface ranges from about 1 to 12 mN/m between the softest and stiffest formulations, thus varying by about one order of magnitude (**Fig. 2E**). In addition, micropipette aspiration (38) measurements on single functionalized or non-functionalized droplets show that the effect of the phospholipids adsorption at the oil/water interfacial is negligible (**Tables S2** and **S3**).

The mechanical properties of complex materials are characterized by both an elastic and viscous behaviors (39) that depends on the timescale of the solicitation. We used a microplate rheometer to measure the complex dynamic modulus G*(f) = G’ + *i*G″ of single droplets over a frequency range of biological relevance (40), and to correlate the droplet interfacial tension γ to their storage modulus G’ (34, 41) (**Fig. 2F**).

The experiment consists in trapping a spherical deformable object between two glass microplates, a rigid and a flexible one, and applying a sinusoidal deformation while measuring the resulting amplitude and phase shift at the tip of the flexible microplate. From the applied oscillatory normal stress and the sinusoidal strain (deformation) of the object, one can infer its complex dynamic modulus G*(f) = G’ + *i*G″, with the real part G’ representing the storage modulus (elastic-like response), and the imaginary part G″ accounting for energy dissipation (viscous-like response).

Sinking droplets (Lipiodol, oil denser than water, used here as reference) exhibited a Kelvin-Voigt behavior, with a constant elastic modulus G’, and a viscous modulus G″ proportional to the frequency (**Fig. 2G**) where elastic-like response is dominant (G’ ≫ G″, **Supplementary Note 3**). We found that, for a fixed strain (*ϵ* = 0.2), the droplet storage modulus decreases as the inverse of the diameter of the resting droplet (**Fig. 2H**), in agreement with the assumption that the only restoring force resisting compression originates from Laplace pressure ΔP = 4*γ/L*_0_ (**Supplementary Note 3**), with *L*_0_ the resting droplet diameter. Thus, by fitting the storage modulus G’ as function of the droplet diameter, one gets a calibration curve for which we extract an estimation of the droplet surface tension, compatible with the pendant drop measurements (*γ* = 1.21 ± 0.04 mN/m - **Supplementary Note 3, Fig. 2E, Table S1-S2**). Therefore, according to this quantification, we conclude that the droplets used in our experiments, i.e. diameter of 11 μm and tension ranging from 1 to 12 mN/m, exhibit an apparent rigidity (storage modulus G’) ranging from 4 to 30 kPa. As comparison, we used the same method to measure B cell stiffness and found G* = 165 Pa (**Supplementary Note 4** and **Fig. S5**). Antigen presenting cells range from 1 kPa (macrophages (34)) to 5-10 kPa (follicular dendritic cells (15), that are essentially fibroblasts). This indicates that the droplets are, as APCs, several times stiffer than B cells and definitely in the range of the substrate rigidity capable of eliciting a strong B cell signal according to (6).

### B cells actively gather antigens at the synapse

We assessed the impact of antigen enrichment during the synapse formation by using two droplet types, one favoring and the other one hindering such enrichment. We thus used two alternative methods to graft the lipids on the droplets: via the surface where lipids are inserted after the emulsification processes (33), or in bulk where lipids are inserted prior to emulsification (23, 26). Using the microfluidics traps described above to immobilize the droplet-B cell contact, we notice that cells interacting with bulk-functionalized droplets gather the antigen to a sustainable central cluster (**Fig. 3A, Movie S1**) in less than 20 min, similarly to what described on planar lipid bilayers (10) or for other cell types (23). By contrast, surface-functionalized droplets do not allow antigen accumulation (**Fig. 3B**), similar to the negative BSA control (**Fig. 3C**). We further quantified the kinetics of antigen accumulation by an index I_*antigen*_ (**Fig. 3D**) that increases for bulk-functionalized antigen-coated droplets as a monotonic exponential saturation curve over 40 min, with a characteristic time of 5 min (**Fig. 3E**, bulk-functionalized antigen-coated droplets). The plateau is directly related to the accumulated antigen concentration: the cluster is 1.5-fold brighter than the initial time point (**Fig. 3E**). We took advantage of the fine calibration of antigen concentration showed above (**Fig. 2E and Supplementary Note 2**) to conclude that B cells are able to gather up to 75 antigens/*μ*m^2^ on droplets in 15 min (plateau, **Fig. 3E**) functionalized with initially 50 antigens/*μ*m^2^ (Antigen^*low*^) (time zero, **Fig. 3E**). We thus named the two types of functionalization - favoring and hindering antigen gathering - respectively cluster÷ and cluster^-^. We will further inquire whether antigen accumulation is an important feature impacting the polarization of B cells.

**Fig. 3.**
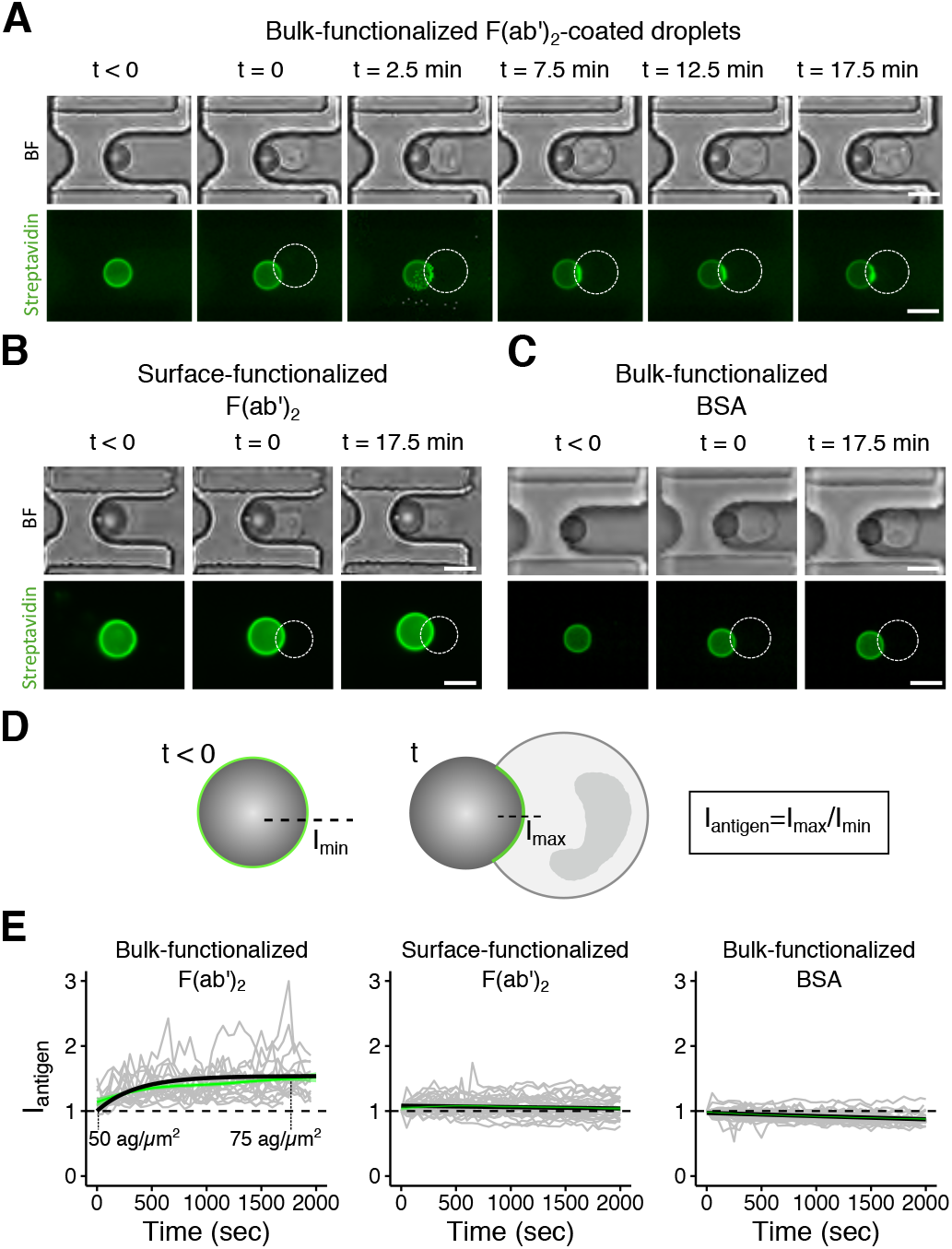
B cells actively gather antigens at the synapse. (A) Time lapse images of the antigen aggregation in a bulk-functionalized droplet-cell contact. All droplets are functionalized with 50 F(ab’)_2_ fragments/*μ*m^2^ on average. Scale bar: 10 *μ*m. (B) Time lapse images of the antigen aggregation (not occurring) in a surface-functionalized droplet-cell contact. Droplets are functionalized with 100 F(ab’)_2_ fragments/*μ*m^2^. Scale bar: 10 *μ*m. (C) Time lapse images of the antigen aggregation in a BSA bulk-functionalized droplet-cell contact. Droplets are functionalized with 100 BSA/*μ*m^2^. Scale bar: 10 *μ*m. (D) Scheme of antigen accumulation analysis. The antigen recruitment index (I_*antigen*_) is defined as the ratio of the intensity at the time zero (I_*min*_) and the intensity over time (I_*max*_), both at the synapse area. (E) I_*antigen*_ time evolution for three different conditions. Only bulk-functionalized antigen-coated droplets show antigen recruitment (left, N=18 cells), while neither bulk-functionalized BSA-coated (right, N=27 cells) nor surface-functionalized antigen-coated droplets (center, N=34 cells) show antigen accumulation. For the bulk-functionalized antigen-coated droplets, the kinetic of antigen accumulation follows *y* = (*p* – 1)(1 – exp(-*t/τ*))+ 1, where *τ* = 5.07 ± 0.14 s. The related plateau p=1.50 ± 0.06 allow to calculate the number of antigens aggregated at the synapse by the cell: drop lets are initially coated with 50 antigens per *μ*m^2^ and cells cluster up to 1.5× 50 = 75 antigens/*μ*m^2^. Green solid curves represent average values of all cells at each time point, and shaded surrounded curves the related 95% confidence interval.

### B cells exhibit different polarization responses after being activated

As a proxy for cell polarization, we followed lysosomes distribution in time - as it has been already intensively studied as a polarization readout (12, 13, 15, 42) - by imaging them using an acidic compartment dye (Lysotracker^®^) over 40 min. This method is fast, reliable and does not interfere with organelles and cytoskeleton dynamics (13). We observe three distinct behaviors for B cells even in contact with the same droplet type (**Fig. 4A**): within 40 min from the contact, lysosomes either stay uniformly distributed in the cytoplasm (no *polarization*), move toward the synapse area (*polarization*, **Movie S2**), or accumulate in the side opposite to synapse (*anti-polarization*, **Movie S3**). We quantified the asymmetry in lysosomes distribution (polarity) by defining a polarization index I_*pol*_ as the ratio between the fluorescence signals in the half cell close and opposite to the droplet, as sketched in **Figure 4B**.

**Fig. 4.**
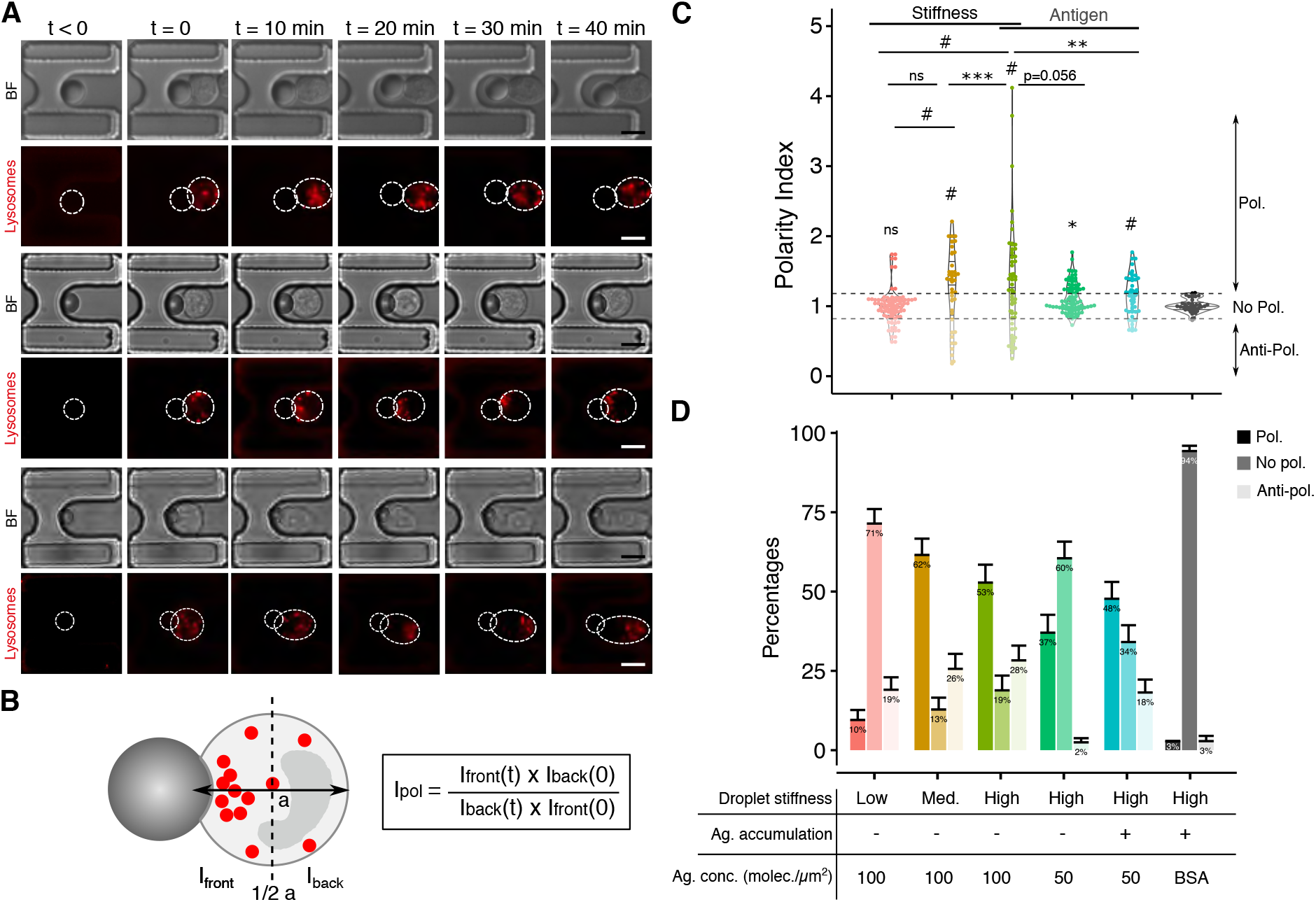
B cells exhibit different polarization responses after being activated, and the onset of polarity is determined by stiffness and local concentration at the synapse. (A) Representative time-lapse imaging of the three cell behaviors: non-polarizing (top, droplet: 4 kPa - cluster^-^ - 100 ag/*μ*m^2^, I_*pol*_ = 0.92), polarizing (center, droplet: 29.8 kPa - cluster^+^ - 50 ag/*μ*m^2^, I_*pol*_ = 1.21), and anti-polarizing (bottom, droplet: 29.8 kPa kPa - cluster^-^ −100 ag/*μ*m^2^ - I_*pol*_ = 0.34) cells. Scale bar: 10 *μ*m. (B) Schematic representation of the polarization analysis and quantification: the polarization index (I_*pol*_) is the ratio between the fluorescence intensity integrated over the front part, in contact with the droplet, and the fluorescence intensity integrated over the back of the cell. The ratio is normalized by its value at time zero. (C) Distribution of the I*_pol_*, *p*-values are computed by pairwise Kolmogorov-Smirnov test –– when not written, the pair is not significantly different. I*_pol_* is the value for which the polarization behavior is sustained, i.e. at 40 min. (D) Percentages of the three cell behaviors depending on the droplet types, error bars are computed as half standard deviation of the percentage obtained by random subsampling 1000 groups of 15 values (typical size of the experimental pool in one day). Stiffness^*low*^, Stiffness^*medium*^ and Stiffness^*high*^ in the text, denotes quantitative stiffness of respectively 4.0, 11.7 and 29.8 kPa. Antigen concentration relative to 100 and 50 antigens/*μ*m^2^ denotes respectively Antigen^*high*^ and Antigen^*low*^. Respective number of analyzed cells from left (pink plot) to right (gray plot): N=50, N=39, N=53, N=85, N=56, N=34, from at least 3 independent experiments. p-values: #: p<0.0001, *** p<0.001, ** p<0.01, * p<0.05.

We investigated the impact of antigen enrichment, antigen concentration and droplet stiffness on lysosome polarization. We compared the cellular responses upon interaction with five different droplets: stiffness^*low*^ (*γ* = 1.7 mN/m), stiffness^*medium*^ (*γ* = 4.7 mN/m) or stiffness^*high*^ (*γ* = 12 mN/m), covered with 50 (Antigen^*low*^) or 100 (Antigen^*high*^) antigen/*μ*m^2^, with bulk- (cluster^+^) or surface-functionalized (cluster^-^) droplets (**Fig. 4C** and **4D**). The panel of activating droplets allows to investigate the effect on polarization of different stiffness (at constant antigen concentration) and different antigen coating conditions (at constant stiffness).

To establish the value of I_*pol*_ for which we consider a cell to be polarized or not, we observe the I_*pol*_ distribution for cells in contact with the non-activating droplet (BSA-coated, gray plot **Fig. 4C**). As spontaneous polarization has been reported (43), we considered two sigmas of the I_*pol*_ distribution for BSA-coated droplets as a threshold. This results in classifying the cells with I_*pol*_ ≥ 1 + 2*σ* =1.18 as polarized. Symmetrically, we defined cells with I_*pol*_ < 1 — 2*σ* =0.82 as anti-polarized and, consequently, 0.82≤I_*pol*_ <1.18 for non-polarized cells (**Fig. 4C**). Furthermore, to check that the polarization occurs specifically upon BCR engagement, we performed two major controls: (i) we excluded an impact of the flow by checking that no cell polarized in absence of interaction at maximal flow of 1.6 mm/s (Control #1, **Fig. S2**); (ii) we excluded an activation from potential soluble antigens coming from the droplet functionalization by observing no polarized cells among the non-interacting ones in the presence of antigens-coated droplet in nearby traps (Control #2, **Fig. S2**). Altogether, these results show that B cells polarize only when they form a specific contact with antigen coated-droplets suggesting that polarization is triggered by specific BCR engagement in our system, and that flow conditions do not alter polarization in our experiments.

**Figure 4C** shows the polarity index distributions of the global cell populations for each droplet condition. For more accurate analysis in each cell population, we classified the cells in the three categories of phenotypes described above (**Fig. 4D**).

### B cell polarization onset depends on antigen concentration and droplet stiffness

We first studied the mechanosensitivity of the polarization process by changing the surface tension of the droplets, hence their effective stiffness. For soft droplets, cells behave essentially as the negative control, although antipolarized cell number increases. However, increasing stiffness from 4 kPa to 11.7 kPa, induces a greater number of polarized cells and with much higher indexes, corresponding to an increased lysosome concentration in the synaptic area. Polarity indexes distributions do not significantly change between stiffness^*medium*^ (4 kPa) and stiffness^*high*^ (≈ 12 kPa). These results show that polarization is a mechanosensitive process and that the threshold of effective stiffness triggering the polarization is between 4 and 12 kPa. Of note, antigen concentration does not depend on droplet stiffness (**Fig. S6**) The major difference between bulk and surface functionalized droplets, antigen gathering, is related to the different fluidity of the surfaces, as we showed by FRAP experiments (**Supplementary Note 5** and **Fig. S7**). Of note, surface tension does not depend on functionalization method (**Fig. S8**). We characterized, therefore, the effects of both antigen concentration and gathering on B cell polarization, by using droplets of similar stiffness (high). Surprisingly, despite not showing antigen accumulation (**Fig. 3B**), cluster^-^ droplets stimulate rather well B cell polarization regardless of antigen concentration. Remarkably, cluster^+^/Antigen^*low*^ droplets, inducing antigen enrichment, are able to generate the same amount of polarized cells (52%) as cluster^-^/Antigen^*high*^ (55%) conditions which initially bear twice as much antigens (respectively 50 and 100 antigens/*μ*m^2^). However, even in polarized cells, polarity indexes are lower in cluster^+^ cells, suggesting that a higher local concentration of antigen stimulates stronger lysosome polarization. In addition, on cluster^-^ droplets, when antigen concentration is halved, the percentage of polarized cells decreases from 55% to 35% (**Fig. 4D**). These results show that by increasing local concentration at the synapse, antigen accumulation overcomes the lack of initial presented antigens and lowers the threshold to established polarity when the density of presented antigens is below 75 antigens/*μ*m^2^.

A recapitulation of all conditions and outcome of polarization is shown in **Table 1**. **Figure 4D** shows that the population of antipolarized cells does not change with stiffness for droplets cluster^-^ coated with a high concentration of ligands, suggesting that for such interfacial conditions, antipolarization does not depend on the mechanics of the lipid droplet. Considering the set of stiff droplets with contrasting interfacial properties, our result show that cluster^+^ droplets with a low ligand level share a similar antipolarization value than cluster^+^ droplets with a high ligand level. On the contrary to the polarization measurement, antipolarization fully disappears for cluster^-^ droplets coated with a low amount of antigens. Although origin of such discrepancy remains elusive, these results suggest that polarization and antipolarization behaviors come from two distinct mechanisms.

**Table 1.** Recapitulation of droplet conditions, properties and outcomes.

| Droplet condition | Oil phase | Surf.tension (mN/m) [pendant drop] | Stiffness (kPa) | Functionalization type | Observed Ag. enrichment | Antigen concentration | Droplet size (mean±sd) (μm) | Polarization | No polarization | Anti-polarization |
| --- | --- | --- | --- | --- | --- | --- | --- | --- | --- | --- |
| (i) | Mineral oil<br>Oleic acid 5%v/v | 1.7 | 4.0 | Surface | No | 100 | 11.06 ± 0.88 | + | +++ | + |
| (ii) | Mineral oil | 4.7 | 11.7 | Surface | No | 100 | 10.04 ± 1.92 | +++ | + | ++ |
| (iii) | Soybean oil | 12 | 29.8 | Surface | No | 100 | 11.0 ± 0.7 | +++ | + | ++ |
| (iv) | Soybean oil | 12 | 29.8 | Surface | No | 50 | 11.0 ± 0.7 | + | ++ | + |
| (v) | Soybean oil | 12 | 29.8 | Bulk | Yes | 50 | 11.30 ± 2.88 | ++ | + | ++ |
| Control - BSA | Soybean oil | 12 | 29.8 | Bulk | No | - | 11.30 ± 2.88 | - | +++ | - |

### The polarization kinetic is affected by droplet mechanical properties

We then assessed the impact of droplet properties on the B cell polarization dynamics by measuring the evolution of the polarization index I_*pol*_ over time (**Fig. 5 A-C**, **Fig. S9** and **Fig. S10**). The kinetics of *polarization (anti-polarization*) events shows increasing (decreasing) curves that reach a steady state about 15 min after the droplet-cell encounter. Lysosomes polarity depends on microtubule network orientation and organelles motility on this network. One single parameter cannot capture this complex kinetics. We chose the model that appears to us by visual inspection of the kinetic curves: an exponential relaxation of the form 1 — exp(–*t/τ*) (or for antipolarizing cells exp(–*t/τ*). By contrast, curves related to *no polarization* show no evolution over 40 min and are centered around I_*pol*_ = 1.

**Fig. 5.**
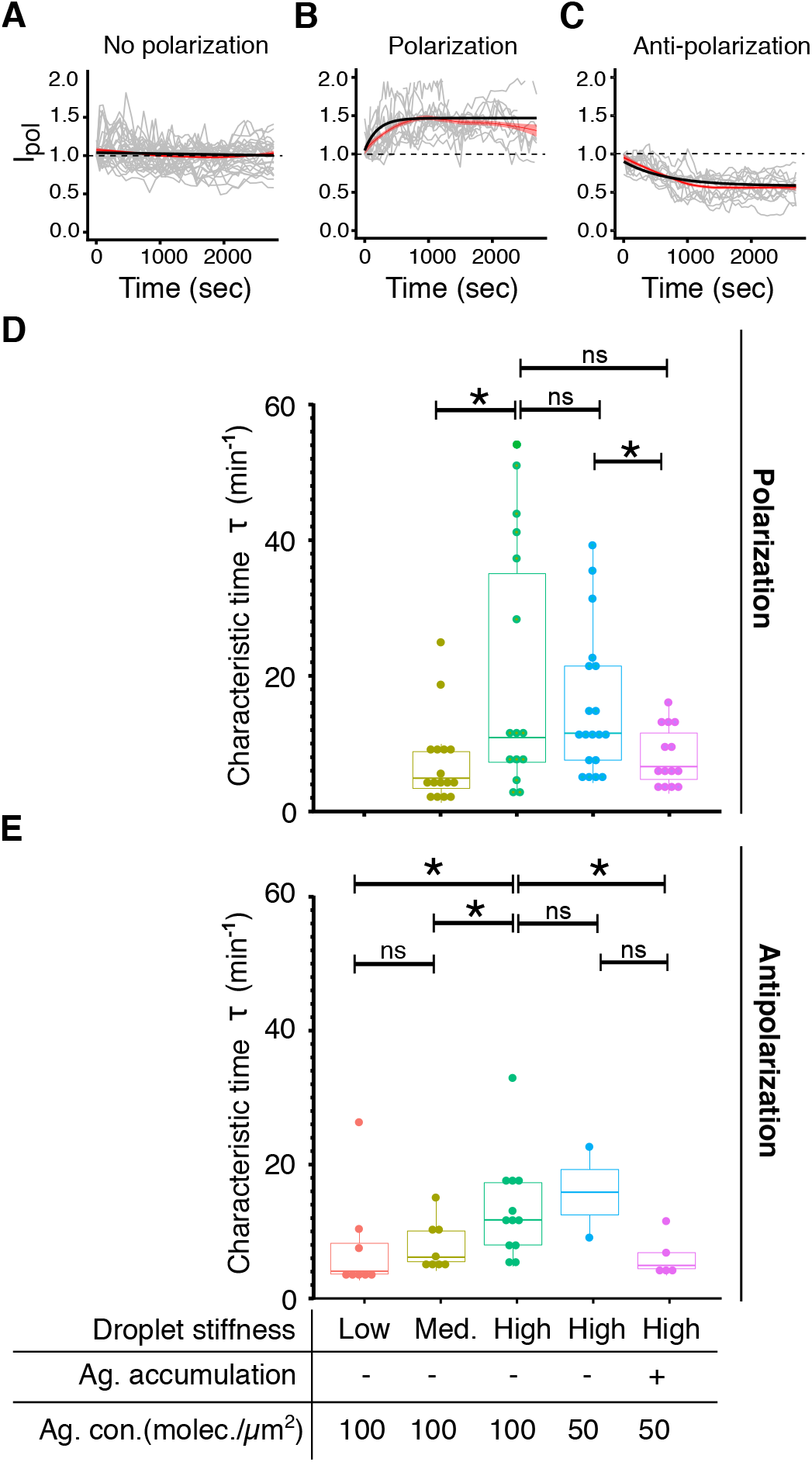
B cell polarization kinetics depends on the mechanical properties of the droplets. (A-B-C) Kinetics of polarization in the three types of behavior for the (A) non-polarized Stiffness^*low*^/cluster^-^/Antigen^*high*^ condition (N=28 cells), polarized Stiffness^*medium*^/cluster^-^/Antigen^*high*^ condition (N=16), and anti-polarized Stiffness^*high*^/cluster^-^/Antigen^*high*^ condition (N=13) cells (from at least 3 independent experiments). Fitting curves of *I_poi_* (black solid curves) for polarized cells and anti-polarized cells respectively follow *y_poi_* = (*p* – 1)(1 – exp(–*t/τ*) + 1 and *y_antipoi_* = (1 – *p*)(1 ‒ exp(–*t/τ*)) + 1 (with *τ* characteristic times of polarization/anti-polarization kinetics). Red shaded curves represents the average values of dynamics at each time point and their 95% confidence interval. (D) Characteristic times of polarization and (E) anti-polarization kinetics (Mann-Whitney tests). Stiffness^*low*^, Stiffness^*medium*^ and Stiffness^*high*^ denotes quantitative stiffness of respectively 4.0, 11.7 and 29.8 kPa. Antigen concentration relative to 100 and 50 antigens/*μ*m^2^ denotes respectively Antigen^*high*^ and Antigen^*low*^. p-values: * p<0.05 and ns: not significant.

From the plateauing exponential curves of *polarization* and *anti-polarization* events (**Fig. 5 B-C**), one can extract the characteristic times *τ* of the respective kinetics (**Fig. 5D** and **E**). This time is inversely related to the rate of polarization, i.e. larger values of *τ* indicate slower kinetics. Figure 5D shows similar characteristic times for cluster^-^/Antigen^*low*^ and cluster^-^/Antigen^*high*^ droplets, meaning that, independently of the antigen concentration, once the polarization is triggered, it reaches a plateau within 15 min (mean value). This suggests that polarization kinetics do not depend on the initial antigen concentration, as long as these conditions are sufficient to initiate polarization. By contrast, characteristic times are significantly different for stiffer versus softer droplets, suggesting that stiffness affects the polarization dynamics: B cells polarize faster in the case for a stiffness of about 12 kPa than 30 kPa. Noteworthy, polarization is faster when droplets allow accumulation of antigens at the synapse (at equal conditions, see stiffness^*high*^/cluster^+^/Antigen^*low*^ vs stiffness^*high*^/cluster^-^/Antigen^*high*^). Although the differences in some cases are not significant (most likely due to low statistics of anti-polarization cell subpopulations), similar tendency is found in antipolarizing behavior.

Altogether, these results reveal that B cells are dynamically mechano-sensitive, as they are able to discriminate stiffness changes, and significantly adjust the polarization rate according to the apparent stiffness of the substrates they face. Also, accumulating antigens at the synapse allow cells to polarize as fast as on softer (stiffness^*medium*^) substrates. Therefore, the local antigen concentration is an important feature to induce an efficient polarization, suggesting a synergistic role of low stiffness and antigen mobility in accelerating lysosome polarization.

## Discussion

In this work we introduced a new material to activate B cell: lipid-coated droplets. We characterized the physical and chemical properties of functionalized droplets and studied their effect on lysosome polarization in B cells. We could prove that these objects are able to activate B cells and that the onset of polarization is sensitive to ligand concentration and droplet apparent stiffness. This is consistent with previous observations showing that lysosome polarization, necessary for protease secretion, antigen degradation and extraction (13), occurs predominantly on non-deformable/stiff substrates, where mechanical internalization is not possible (15). We could not access whether there is antigen internalization from the droplets. However, in a recent work we show that there is, at least on stiff droplets, exocysts enrichment at the synapse, a signature of proteases release (44). Future efforts will be directed to make droplets where mechanical extraction is possible.

We showed that B cells polarize only when they interact with substrates presenting a surface tension greater than 4.7 mN/m (corresponding to an apparent stiffness of 11.7 kPa). This is consistent with observation on macrophages and follicular dendritic cells (15). Of note, the stiffest (11.7 and 29.8 kPa) and softest (4.0 kPa) droplets induce responses similar to, respectively, follicular dendritic cells and dendritic cells (15). We propose that this is linked to the BCR mechanosensitivity. Indeed, BCR signal is known to depend on the substrate stiffness (6) and the signal downstream the BCR to induce the accumulation of polarity cues (3, 12, 16). These polarity cues establish the pole by anchoring the centrosome (45), reorienting the microtubule based traffic towards the synapse: the stronger the signal, the stronger the establishment of polarity cues, the higher the probability of polarizing. Unexpectedly, we also found that the kinetic of polarization is sensitive to the model APC stiffness. Mechanosensitivity of dynamical properties has been shown in adhesive cells: actin flows increase with the rigidity of the substrate (46), cells adjust their contractility in real time to the stiffness change of the substrate (47), and T cells adapt their pulling force at the synapse to the stiffness of the interacting object (48, 49). One explanation for the dependency of polarization kinetics on the mechanics of the substrate, could be that the increase in stiffness downstream of the BCR activation (see e.g. (50)), hinders the displacement of the centrosome and hence the polarization of lysosomes, effectively lowering the polarization rate on the stiffest substrates. Although we have no direct readout of the signal, it is interesting to compare this results with recently published data on cytotoxic granules release in CD8+ T cells (4): in this case the stronger the signal, the highest the frequency of polarizing cells. However, the kinetics of polarization does not seem to be influenced by the strength of the signal. Future investigations on the relation between signal, rheological changes and polarization following BCR activation will better highlight differences and analogies between the two lymphocytes.

We quantified the statistics of cell polarization when they were able to accumulate antigens at the contact zone or not, and to interact with 50 to 100 antigens/*μ*m^2^. Our results showed that the maximal amount of polarized cells is reached for 75 antigens/*μ*m^2^, suggesting that above this density, antigen concentration has no effect on the polarization process. This value is compatible with the maximal density of BCRs available in an activated synapse^1^. In addition, previous work reported that antigen concentration from 15 to 150 antigens/*μ*m^2^ modulates the B cell activation with a threshold that depends on the affinity (10). Although we cannot quantitatively compare these results to ours (different antigens and presentation assays), this suggests the concentration threshold for polarization might also depend on the antigen affinity. Further experiments will better elucidate the link between early signaling, lysosome polarization, and antigen affinity. Other works have shown that B cells interacting with mobile ligands displayed significantly greater signaling because of the formation, the accumulation, and the merging of BCRs microclusters (11, 42, 52). We herein show that lysosomes polarize independently of the ability to form a central antigen cluster. We showed, however, that “mobility”, allowing the local increase of antigen concentration at the synapse, helps B cells overcome the low initial antigen concentration (below 75 antigens/*μ*m^2^), increases the chance for a cell to polarize and its polarization rate. This suggests that local antigen concentration is at the synapse is a crucial feature for inducing polarization and make it quicker.

One interesting aspect of our system is to allow stimulation of B cells using the same presentation assay, that combines a fluid (such as bilayers) and soft material (such as gels) to investigate new synergies. In addition, the droplets are a highly reproducible and well-calibrated tool in terms of chemical and mechanical properties (compared e.g. to beads or gels), which makes them adapted to study cell-to-cell variability. This has allowed us to show that in clonal B cell lines, not all cells polarize their lysosome towards the synapse, but some even displace them in the opposite side of the synapse (anti-polarize). In T lymphocytes, a transient anti-polarization of actin and myosin has been observed in the early stage of the T cell/APC contact, and does not depend on antigen recognition (53). By contrast, B cell anti-polarization is stable (over 40 min), BCR specific and is only triggered above 75 antigens/*μ*m^2^. Interestingly, the stiffness does not influence the onset of anti-polarization. Like the polarization process, antigen mobility and concentration does not affect the anti-polarization kinetics, but stiffness does. Of note, we observed that anti-polarized cells are more elongated and considerably less round than the polarizing or non responsive ones. Similar events have been described in frustrated T cell-APC conjugates (54), where the centrosome is blocked behind the nucleus while membranes accumulate at the synapse, which ultimately deforms the cell. Alternatively, the anti-synapse might represent a signaling complex in its own right as already described in T cells (53). Further comparison of the proteins implicated in the proximal and distal poles, both during the polarization and the anti-polarization formation, would elucidate this latter suggestion.

Compared to existent antigen presenting surfaces, our method allows us to modify fluidity, concentration and stiffness on the same material. While we achieved similar results as on bilayers (for concentration/ fluidity) and rigidity threshold for activation, we will investigate more thoroughly (e.g. looking at signal magnitude) the interplay between the different properties in a future work. Our method allowed us to precisely phenotype lysosome polarization dynamics. This may be combined with classic (genetic or pharmacological) perturbations to dissect single cell polarization mechanisms in real time and further understand the biological significance and functions of the phenotypes we described. We anticipate that this approach will be of interest in many other contexts, where cell-cell variability is important for the immune function such as T cell immune synapses (55), cytotoxic cell encounter (56, 57), and phagocytosis (23, 58).

## Materials and methods

### Materials

Pluronic F-68 (Poloxamer 188, CAS no. 9003-11-6), sodium alginate (CAS no. 9005-38-3), Tween 20 (polyethylene glycol sorbitan monolaurate, CAS no. 9005-64-5), oleic acid (CAS no. 112-80-1, ref. O1008), silicone oil (viscosity 350 cSt at 25°C, CAS no. 63148-62-9) and mineral oil (light oil, CAS no. 8042-47-5) were purchased from Sigma-Aldrich (Saint Quentin Fallavier, France). DSPE-PEG_2000_-Biotin lipids (1,2 - distearoyl - sn - glycero - 3 - phosphoethanolamine - N - [biotinyl (polyethyleneglycol) - 2000] ammonium salt, CAS no. 385437-57-0) were purchased from Avanti Polar Lipids (Alabaster, AL, U.S.A). FluoProbes 488 streptavidin (ref. FP-BA2221) and Biotin-SP (long spacer) AffiniPure F(ab’)_2_ Fragment Goat Anti-Mouse IgG, F(ab’)_2_ fragment specific (min X Hu, Bov, Hrs Sr Prot) (CAS number: 115-066-072) were respectively purchased from Interchim (Montlucon, France) and Jackson ImmunoResearch (Ely, U.K.). Ultrapure water (Millipore, 18.2 M.cm) was used for all experiments. All reagents and materials were used as purchased, without any further purification. Lipiodol (CAS no. 8002-46-8) was kindly provided by the company Guerbet (Villepinte, France).

### Cell culture

Mouse IgG+ B-lymphoma cell lines IIA1.6 (31), derived from A20 cell lines (ATTC# TIB-208), were cultured at 37°C in a 5% CO2 atmosphere in CLICK medium (RPMI 1640 Medium GlutaMAX™ Supplement, ref. 61870036; supplemented with 10%v/v decomplemented fetal calf serum, ref. 16140063; 1%v/v antibiotic Penicillin-Streptomycin, ref. 15070063; 2%v/v sodium pyruvate at 100 mM, ref. 11360070; and 0.1%v/v 2-Mercaptoethanol at 50 mM, ref. 31350010). All these reagents were purchased from Life Technologies - Gibco. Experiments were performed with cell densities close to 106 cells/mL, which were diluted to ca. 200 000 cells/mL and split in 10 samples of 1 mL, inserted in a 12-well plate (Falcon, ref. 353043). For each experiment, a 1 mL cell sample and 25 *μ*L of HEPES (4- (2-hydroxyethyl) - 1- piperazineethanesulfonic acid, Gibco) were put in a 1.8 mL tube compatible with the pressure controller (Fluigent, France). In the meantime, other samples remained incubated at 37°C, 5% of CO2 for a maximum of 4h.

### Cell staining

Lysosomes were stained by loading the cells with LysoTracker™ Red DND-99 (1 mM, ref. L7528) at a final concentration of 50 nM purchased from Life Technologies | Thermo Fisher Scientific (Invitrogen), into CLICK Medium, 37°C, 5% CO_2_ for 30 min. The cell concentration is about 0.5 M cells/mL.

### Bulk droplet functionalization protocol with phospholipids

The lipid-containing oil was obtained by dilution of DSPE-PEG_2000_-Biotin phospholipids (Avanti Lipids, Alabama, USA) in soybean oil at 0.03 mg/mL, followed by 30 min of sonication and evaporation of the chloroform from the oil at room temperature. This oil was dispersed and emulsified by hand in an aqueous continuous phase containing 15% w/w of Pluronic F68 block polymer surfactant and 1% w/w sodium alginate (Ref: W201502, Sigma Aldrich, St. Louis, MO, USA) at a final oil fraction equal to 75% w/w. The rough emulsion was then sheared in a Couette cell apparatus at a controlled shear rate following the method developed by Mason *et al*. (59) to narrow the droplet size distribution to 12.4 *μ*m ± 2.3 *μ*m (**Table 1**). For storage, emulsions are diluted to an oil fraction of 60 wt % with 1 wt % Pluronic F68 in the continuous phase and stored at 12°C in a Peltier storage cupboard for several weeks.

### Surface Droplet Functionalization Protocol with Phospholipids

Droplets were formulated using a Shirasu Porous Glass apparatus (SPG Technology Co., Japan) by extruding the oil phase (with or without 5%v/v of oleic acid) through a ceramic membrane (with pores of 3.1 *μ*m, SPG Technology Co., Ltd) within an aqueous solution containing 15%v/v of Pluronic F68, continuously and vigorously stirred. We obtained droplets measuring 10.98 ± 0.68 *μ*m in diameter. Once stabilized, droplets were washed out 3 times with an aqueous solution of Tween 20 at CMC (0.0007% w/v). The supernatant was removed and replenished after each centrifugation step (30 sec at 2000 RPM). The suspension is centrifuged and rinsed four times with phosphate buffer (PB)/ Tween 20 to decrease the amount of Pluronic F68 in the continuous phase. After the last rinsing step, most of the continuous phase is removed from the microtube to decrease the total volume of emulsion to 10 *μ*L. We then add 10%v/v of the DMSO stock solvent solution containing the phospholipids. We then add a phosphate buffer supplemented with Tween 20 at the CMC (PB - Tween 20) to reach a total volume of 200 *μ*L of suspension. Droplets are incubated for 30 min at room temperature in the presence of lipids in the bulk phase and finally rinsed several times with a PB - Tween 20 buffer to remove the phospholipids in excess. The quantity of phospholipids available in the bulk phase at the initial stage is adjusted by diluting the stock solutions in DMSO so that the volume fraction of co-solvent remains constant for all experimental conditions. For a given working volume (200 *μ*L), the lipid concentration in the bulk phase is expressed as a droplet surface area equivalent. One equivalent (eq.) corresponds to the number of molecules required to cover the total area of a droplet sample with a compact monolayer (33).

### Coating of Biotinylated Droplets with Streptavidin and F(ab’)_2_ fragments

After the first step of emulsification and insertion of biotinylated lipids at the interface, 1 equivalent of streptavidin solution at 1 mg/mL was added, followed by an incubation time of 30 min and a washing step to remove all excess streptavidin. Finally, 1 equivalent of F(ab’)_2_ fragments (1 mg/mL) or BSA (1 mg/mL) was added to the droplet solution, followed by an incubation time of 30 min and a washing step to remove all excess F(ab’)_2_ or BSA. One equivalent (1 eq.) of a given macromolecule (lipid or protein) corresponds to the theoretical number of molecules needed to cover the entire surface with a monolayer of molecules or proteins, considering the available surface of droplet and molecule dimensions as previously reported in (33).

### Measurement of the adsorption isotherms of the surface-functionalized droplets

To quantify the amount of antigens presented by droplets, we first titrated the antigens (from 0.05 to 5 eq.) bound to streptavidin molecules (1 eq.) and biotinylated DSPE-PEG_2000_ lipids (100 eq.). We revealed the antigens coating with a Donkey anti-Goat IgG (H+L) Cross-Adsorbed Secondary Antibody (Alexa Fluor 647, Invitrogen, ref. A-21447) and analyzed the fluorescence intensity. In our case, the droplets are saturated for 4 eq. of F(ab’)_2_ fragments (**Supplementary Note 2**). We then converted this relative concentration into absolute value by comparing fluorescence intensity of droplets and commercial tagged beads (Quantum™ MESF, Bang Laboratories, Inc.). We assessed the related fluorescence intensities of droplets coated with a concentration of streptavidin ranging from 0 to 2 eq. and saturated in biotinylated DSPE-PEG_2000_ lipids (100 eq.) (**Supplementary Note 2**). These values are added on a calibration curve correlating the molecular equivalent surface fluorescence (MESF) of beads (Quantum™ MESF, Bang Laboratories, Inc.) with their fluorescence intensities (**Supplementary Note 2** and **Fig. S4**). Finally, we converted the droplet MESF into absolute values of F(ab’)_2_ fragments in antigens per *μm^2^* (**Fig. 2D**) depending on different droplet and protein features. We obtained that droplets present from 0 to 150 antigens/*μ*m^2^ when they are functionalized with DSPE-PEG_2000_-Biotin (100 eq.), streptavidin from 0 to 2 eq. and F(ab’)_2_ fragments (4 eq.). For the rest of the experiment, we use droplets coated with 50 and 100 antigens/*μ*m^2^. Fluorescence measurements in **Fig. S6** have been performed using Alexa Fluor^®^ 647 Mouse IgG1, κ Isotype Ctrl (ICFC) Antibody, Biolegend 400135, at a concentration of 0.5 eq.

### Measurement of oil/water interfacial tension by the pending drop technique

The pendant drop technique consists in inferring the interfacial tension from the shape profile of a pendant drop of one liquid in another at mechanical equilibrium. We used a pending drop apparatus (drop shape analyzer, Krüss, PSA30) to measure the interfacial tension of the various oils considered in this study - mineral oil, mineral oil supplemented with 5%v/v of oleic acid, soybean oil, lipiodol and lipiodol supplemented with 5%v/v of oleic acid - versus an aqueous solution composed of 15%w/w Pluronic 68. The interfacial tensions reported in **Table S1** correspond to the equilibrium value reached when the drop surface has been entirely covered by surfactants coming from the continuous phase.

### Measurement of droplets interfacial tension by the micropipette technique

The micropipette-aspiration method that has been described in detail in a previous article (38). Micropipettes were made from 1 mm borosilicate glass-tube capillaries (Harvard Apparatus, USA) that were pulled in a pipette heater and puller (P-2000, Sutter instrument Co., USA) to tip diameters in the range of 3 to 5 *μ*m. A 3 axis-micromanipulator (Narishige) allowed for pipette positioning and manipulation. The pipette was connected to a pressure controller (Fluigent) to apply precise negative pressures. A solution of sinking droplets, made with lipiodol oil coated or not with phospholipids, was inserted into a glass/coverslip chamber. The pipette aspired the droplet until reaching an equilibrium where the elongation part is equal to the pipette inner dimension. The interfacial tension *γ* is *γ* = ΔP· (R_*p*_/2) where ΔP is the negative pressure and R_*p*_ the micropipette radius. Upon aspiration by the very thin glass pipette, the droplet deforms and a spherical cap of radius R_*c*_ forms at the tip of the pipette. At equilibrium, the value of R_*c*_ depends on the interfacial tension *γ* of the droplets, the radius of the droplet R_*D*_, the aspiration pressure ΔP and can be expressed as:

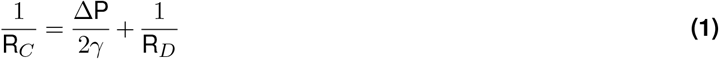

The aspiration ΔP corresponds to the pressure difference between the inside of the pipette and the external pressure. *γ* and *R_p_* are constant throughout the experiment. Hence, varying ΔP induces changes in R_*c*_: the larger ΔP, the smaller *R_c_*. During measurements, ΔP is slowly increased and R_*c*_ decreases until reaching the radius of the pipette R_*p*_. Up to that critical aspiration ΔP_*c*_, little change is observed in the geometry of the system. As soon as ΔP becomes greater than ΔP_*c*_, R_*c*_ becomes smaller than R_*p*_, which results in the sudden entry of the oil droplet in the pipette. This provides a direct measurement of the surface tension of the droplet:

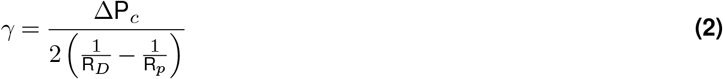

The interfacial tensions reported in **Table S2** have been measured via micropipette on phospholipid-fonctionalized and non-functionalized droplets. Droplets are stabilized by F68 and bath into an aqueous phase made of phosphate buffer/Tween 20 at CMC.

### Measurement of droplet mechanical properties by the microplate rheology

The microplate micro-rehology experiment consists in applying a sinusoidal displacement to the base of a flexible microplate B(*t*) =B_0_*e*^*iωt*^ (with elastic constant *k*) and measuring the resulting amplitude T_0_ and the phase shift *ϕ* of the flexible microplate tip in contact with the droplet (see **Fig. 2F**). Of note, the offset of the microplate applies a strain of 2 μm and the oscillation amplitude of 1 μm is applied around this point. The droplet complex dynamic modulus G* is obtained from the applied oscillatory normal stress. Microplates were made from glass lamellae of 100 x 2mm with 0.1-0.3 mm thickness. The rigid plate can be either pulled out of 0.2 or 0.3 mm original thickness, but must be kept short to ensure a high stiffness. Lamellae were heated and pulled (Narishige PB-7, Japan) until breaking in two similar parts. Microplates were calibrated using a microplate of known bending stiffness as a reference. The bending stiffness of the reference microplate has been initially determined by stacking copper micro-wires at the extremity of the plate, following the protocol reported in (60). A microplate of low bending stiffness (flexible) acts as a spring, and a rigid plate about 1000 times stiffer than the flexible one acts as support. During experiments, the flexible plate is oscillated, applying oscillating stress on the studied sample and compressing it sinusoidally against the rigid plate. The flexible plate was oscillated from 0.1 to 6.4Hz for droplets and until 1.6 Hz for cells. Micrometric displacements of plates are controlled via highly-resolutive piezoelectric micromanipulators. The deflection *δ* of the flexible plate is related to the force *F* exerted on the sample by *F* = *kδ* where *k* is the bending stiffness of the flexible plate. The deformation of the sample is *ϵ* = (*L*_0_ — *L*)/*L*_0_ where *L*_0_ is the resting cell or droplet diameter. For mechanical characterization, we used the dynamic analysis protocol consisting in applying a controlled sinusoidal stress or strain at different frequencies on the cell, while measuring the complementary visco-elastic response provided by the droplet or cell (60). For droplet experiments, Lipiodol (Guerbet) oil (that sediment being denser than water) was used to fabricate droplets and glass microplates were coated with PLL at 0.1%v/v. For cells, only the chamber was coated with Sigmacote (Sigma-Aldrich).

### Measurement of diffusion coefficient - FRAP experiments

Droplets have been coated with DSPE-PEG-Biotin/streptavidin and F(ab)’_2_ and then image with a spinning-disk confocal microscope Nikon Eclipse 2 equipped with a Yokogawa CSU head, objective Apochromat 63X NA1.3. Droplets were photobleached with circular ROIs of D = 2.5, 5.1 and 8.8 *μ*m at the top of their surface following (61). Movies of the recovery were taken with a frame rate of 30 s before and after a photobleaching pulse of 150 ms. Experiments were performed at 37°C. The recovery time was inferred from exponential fit of the measured intensity. The diffusion coefficient was extracted from a linear fit of the mean of the measured times at different radius (61).

### Design and microfabrication of the microfluidic trapping array

Microfluidic chips were designed on CleWin (WieWeb Software). The double layer technique required the creation of two masks: traps and pillars. Chrome masks were fabricated with the *μ*PG 101 maskless aligner (Heidelberg Instruments Mikrotechnik GmbH), and the final silicon mold with the MJB4 aligner system (Karl Süss). Two SU8 photoresists (Microchem) have been used, 2005 and 2010 to respectively obtain 5 *μ*m-thick pillars and 10 *μ*m-thick traps. To avoid any PDMS sticking to the small SU8 structures and long-term mold damage, we coated the wafer with a fluorinated silane by vapor deposition (trichloro (1H,1H,2H,2H-perfluorooctyl) silane, CAS: 102488-49-3, Merck-Sigma-Aldrich). Liquid PDMS (RTV 615, Momentive Performance) at a base:crosslinker 1:10 ratio was poured on the silicon-SU8 mold and cured for more than 2h at 70°C to complete the crosslinking. After cutting PDMS pieces and punching out inlets and outlets with a biopsy puncher (OD = 0.75 mm, Electron Microscopy Sciences, Hatfield, UK), the top PDMS part was bonded to a glass-bottom Petri dish (FluoroDish FD35-100, WPI) together after a O_2_ plasma treatment of both surfaces (50 W for 30 s, 20 sccm O_2_ flow, 0.15 torr pressure, Cute Plasma oven, Femto Science, Korea), and left for 30 sec at 90°C to improve the bonding. For a long-term hydrophilic coating (Hemmila, 2012) of the inner PDMS channels, a 0.25 wt% polyvinylpyrrolidone (PVP, K90, Sigma Aldrich) solution was injected in the chip. Finally, the chip channels were rinsed with culture media before inserting droplets and cells.

### Experimental setup for the cell-droplet encounters

By connecting the microfluidic devices to a pressure controller, a fixed pressure drop ΔP between the inlet and the outlet of the chamber is set (maximum 1000 Pa, corresponding to a fluid velocity of 1.6 mm/s). Traps are initially rinsed with CLICK medium during at least 5 min. Then, 200 *μ*L of the droplets suspension (10^6^ droplets/mL in CLICK medium) are inserted in the microchip using the pressure regulator. Once a desired number of droplets is trapped, the first tubing is carefully removed and replaced by the tubing connected to the B cell-containing tube. Different positions and focus are marked and cells are progressively inserted at a maximum speed of 1.6 mm/s while the acquisition is launched. To avoid a saturation of traps in cells while ensuring culture medium replenishment, the flow is lessened at 0.4 mm/s. Particle displacements and trapping are observed by video-microscopy.

### Microscopy-Imaging of B cell polarization dynamics

Brightfield and fluorescent images of the synapses are acquired on a Leica DMI8 microscope (Germany) connected to an Orca Flash4.0 sCMOS camera (Hamamatsu Photonics, Japan). Epi-illumination is done with a LED light (PE-4000, CoolLED) and a GFP filter set (Excitation/Emission: 470/525 nm) for the fluorescent coated droplets, and a TexasRed filter set (Excitation/Emission: 561/594 nm) for the fluorescent cells. Time zero of the experiment is defined manually when a cell encounters a droplet. All pictures of cell-droplet pairs were imaged with a 40x objective (Leica, dry, N.A.=0.8, Framerate: 50 sec.).

### Image Analysis

Image analyses were performed with ImageJ/ Fiji (62) (version 1.52i), and data analyses were performed with R (RStudio) software (version 2). For all processes, 16-bit images were analyzed. Codes are available on request.

### Quantification of antigen recruitment

The recruitment index is defined as the ratio between the fluorescence intensity over time I(t) divided by the fluorescence intensity at the initial time I(0), both at the synapse area. The synapse area is countered by hand for each time point. The maximum and mean fluorescence intensities were measured at each time point (from time zero to 40 min).

### Quantification of lysosome polarization and cell classification

Each cell shape was contoured by hand for each time point (from time zero to 40 min) and using a ImageJ macro, automatically divided in two parts according to the main orientation of the cell-droplet pair. It extracted the fluorescence intensity at the middle front side of the cell (in contact with droplet) I*_front_*(*t*) and at the back of the cell I_*back*_(*t*) at each time point, and computed the polarization index defined as [**I**_*front*_(*t*)/**I**_*back*_ (*t*)] × [I_*back*_(0)/I_*front*_(0)] (**Fig. 4D**). To classify the cells we noticed that most of I_*pol*_ curves reach a plateau, hence focused on the final I_*pol*_ defined as averaged over last 5 minutes (in the rare case where this was not possible as the polarization kinetics followed a non-plateauing growth we considered only the last time point.). Transient polarizing cells were not classed as polarized.

### Numerical simulations

The numerical simulations are performed with COMSOL Multiphysics ^®^(version 5.3). We first developed a two-dimensional depth-averaged computational fluid dynamics - finite element model of the microfluidic chamber with exactly the same approach that the one proposed in (63). We consider a planar and incompressible flow, at steady state, and a Newtonian fluid with a dynamic viscosity of *μ* = 10^−3^ Pa.s. Hence, the equation set verified inside the fluid domain is written as -▽P+*μ*Δ*v* + *f_v_* = 0 and ▽ · *v* = 0, where P is the pressure field, *v* the velocity field and *f_v_* the volume force. In addition, the non-slip condition on the out-of-plane walls is considered with a Darcy’s law (also called “shallow channel approximation”) thus follows *f_v_* = −12μ/*d*^2^*v*.

On physical walls, a non-slip boundary condition is added. The pressure is set as uniform both at the inlet and outlet. This pressure is equal to the pressure drop in the chamber or equal to 0 Pa, respectively at the inlet and outlet. Finally, we deduced the pressure drop through a single trap ΔP_*trap*_ from the isobar lines plot (**Supplementary Note 1**).

A 3D model focused on the geometry of one trap. We used the same hypotheses and set of equations than the previously detailed 2D model except that we do not consider Darcy’s law, thus we consider a non-slip condition on the upper and lower faces of the chamber as *f_v_* = 0. On the inlet, the pressure is set as uniform and equal to ΔP_*trap*_ while on the outlet, it is considered as uniform and equal to 0 Pa. Finally, we considered the flow as symmetric on the boundary linking with the rest of the microfluidic chamber.

### Statistical analysis

All statistical analysis have been performed using Prism GraphPad (version 8) and R programming language. Codes are available on request. Tests and p-values are specified in the figures.

## Data availability

ImageJ macro, R code and microfluidic trap designs are available in the following GitHub folder: https://github.com/FattaccioliLab/PhenotypingBCellsWithLipidDroplets.

## ACKNOWLEDGEMENTS

The authors thank A.M Lennon-Duménil for scientific support and critical reading of the manuscript, thank the company Guerbet for having kindly provided Lipiodol^®^oil samples, F. Pincet, D. Langevin, B. Cabane and J. Voldman for helpful discussions, P. Saez for cell culture advice and O. Ait-Mohamed for help with statistical analysis. This work benefited from the technical contribution of the Institut Pierre-Gilles de Gennes joint service unit CNRS UAR 3750. The authors would like to thank the engineers of this unit for their advice during the development of the experiments. The authors are grateful to H. Moreau, J. Brujic, M. Thery and R. Dreyfus for critical reading of the paper and suggestions.

## AUTHOR CONTRIBUTIONS

L.P., P.P. and J.F. designed the study. L.P. performed all the experiments and analyzed experimental results. N.R and R.A. conducted all simulation studies. O.M carried out preliminary experiments on microfluidics and cell culturing. J.P. helped during microfluidics and cell culturing experiments. D.C. made the micropipette setup and contributed to the experiments. A.C. and C.L.G. helped with Imagestream experiments described in the answer to the referees. S.A and A.A performed all microplate experiments, analyzed experimental results and wrote the manuscript. L.P., P.P and J.F. wrote the manuscript. All authors approved the final version of the article.

## Supplementary Information

### Supplementary Note 1: Microfluidic Chip Computational Fluid Dynamics Characterization

The 15 μm-thick U-traps can immobilize 12 μm-large droplets and B lymphocytes (IIA1.6 cell lines) of comparable size. The traps are 10 μm-thick traps and are raised by 5 μm-thick pillars to ensure a continuous flow into the chip (**Fig. 3C**). Thanks to the pillars, the fluid streamlines are not deviated when a first object is trapped in the weir structures, hence they allow the easy capture of a second object such as a B lymphocyte.

We sought to determine the inlet pressure to impose into the chamber to mimic a shear stress in the same order of magnitude of 0.6 Pa, representing the wall shear stress close to the afferent vessel (29). We first modeled the fluid flow into the entire designed microfluidic chamber with a 2D depth averaged Computational Fluid Dynamic (CFD) model according to inlet pressures from 0 to 1000 Pa. We deduced the corresponding pressure gap occurring around each trap thanks to isobar lines (**Fig. 1D** and **1E**). From these pressure gap values we built a 3D CFD model of the flow inside and around a U-trap occupied by a droplet and a B-cell that we considered as rigid bodies. This approach is very similar to the one proposed in (64). From the fluid velocity field, we calculated the fluid shear rate and deduced the fluid shear stress norm applied on the B cell (considering the fluid as Newtonian). Particular attention was paid to ensure mesh convergence of the results using boundary layer elements and mesh refinement on the B-cell boundaries. We obtained a maximal shear stress on B cell between 0.07 and 0.75 Pa (lower blue dashed and pointed curve in **Fig. S3 C-D**) for the least loaded trap and between 0 and 1.0 Pa for the most loaded trap (above red dashed curve **Fig. S3 C-D**). These values suggest that shear stress occurring on the B cell in our device is totally comparable to the in-vivo shear stress experienced by B cells in lymph nodes.

### Supplementary Note 2: Titration of antigen concentration on droplet surface

We first titrate streptavidin attachment by measuring the adsorption isotherm onto the phospholipid-coated droplet surface (see **Materials and Methods** section for details). The bulk concentration of streptavidin is expressed in molar equivalents, i.e. the theoretical amount of proteins necessary to cover the droplet sample with a close packed monolayer as described in (33). For instance, one equivalent of streptavidin for 10 million of 11 *μ*m-large droplets corresponds to 4.10^-10^ moles of proteins (**Table S3**). The fluorescence of the droplets is characterized by epifluorescence microscopy, following the method detailed in Pinon *et al*. (33). The streptavidin adsorption isotherm fits with a Langmuir isotherm with a K_*Strep*_, the streptavidin concentration producing half occupation, equal to 9.7 ± 4.7 eq^-1^ compatible with our previous analyses (**Figure 1D**) (33).

For a bulk concentration of streptavidin corresponding to a maximal surface coverage (1 eq.), we measured the adsorption isotherm of antigens (F(ab’)_2_) using fluorescent secondary antibodies and epifluorescence microscopy for the quantitative fluorescence measurements of the total amount of proteins attached to droplets. **Figure 1E** shows that the antigen adsorption isotherm fits with a Langmuir isotherm with a K_*Ag*_ = 1.74 ± 0.74 eq^-1^. We finally converted the streptavidin fluorescence intensity value in a molecular equivalent of proteins using fluorescence calibration beads, and knowing the average number of dyes attached per streptavidin (and therefore per F(ab’)_2_, **Supplementary Note 2**).

The fitting curves of titrations used in **Fig. 2** follow Langmuir isotherms, hence the fluorescence intensity varies in time as

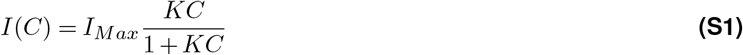

where *C* is the bulk concentration of the molecule of interest, *K* the affinity constant and *I_Max_* the fluorescence intensity at large *C*.

Molecules of equivalent soluble fluorophore (MESF) is converted to antigen surface concentration as follows:

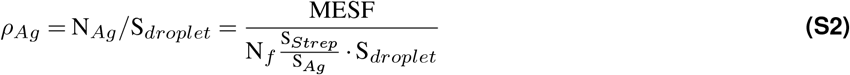

where S_*Strep*_ and S_*Ag*_, are the respective geometric area of a streptavidin molecule (65) and of a F(ab’)_2_ fragment (66), S_*droplet*_ is the droplet surface, and N_*f*_ is the number of fluorophores per streptavidin (as streptavidin is tetravalent, and one bond is occupied by the lipid biotin, we considered N_*f*_=3).

### Supplementary Note 3: Elastic and viscous moduli of oil droplets

Elastic-like behavior is characterized by proportionality between the stress *σ* and strain *ϵ*, typically *σ_e_* = *Eϵ*, with E the elastic modulus characteristic of the material. Viscous response is characterized by proportionality between the stress *σ* and strain rate 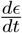, typically 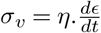, with *η* the viscosity of the material. When elastic and viscous stresses add up, and using complex notation to describe dynamic rheology (oscillating stress and strain at *ω* = 2*πf*, where *f* is the frequency), one gets:

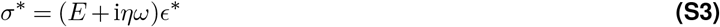

The complex viscoelastic modulus *G*\* is defined as:

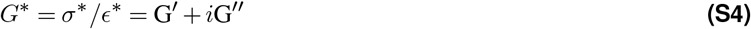

Thus, one finds, for the so called Kelvin-Voigt viscoelastic model:

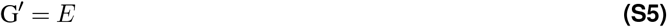

and

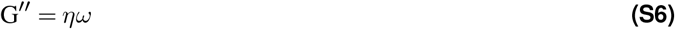

Let us now consider an oil droplet of resting diameter *L*_0_, characterized by a surface tension *γ* and a shear viscosity *η*. When uniaxially compressed between parallel microplates, it develops a restoring (elastic-like) force due to Laplace pressure ΔP = 4 *γ*/L_0_ and a viscous dissipative drag proportional to the elongational viscosity, *η_e_* = 3*η* for a newtonian liquid.

When the droplet is compressed from its resting diameter *L*_0_ to a slightly smaller size *L*, using a Hertz-like model for a small indentation (*L*_0_ — *L*)/2, one finds that the static force resisting compression is simply F_*e*_ = ΔP*πL*_0_ (*L*_0_ — *L*)/4, where *πL*_0_ (*L*_0_ — *L*)/4 is the droplet-microplate contact area and ΔP the Laplace excess pressure. Thus, the elastic-like stress is simply ΔP, and the storage modulus *G*’ = ΔP/*ϵ* (67), where *ϵ* = (*L*_0_ — *L*)/*L*_0_ is the droplet strain. Expressing the modulus as function of the surface tension, one finds G’ = 4*γ*/*ϵL*_0_, hence the fitting curve used in **Fig. 2D** to estimate the surface tension of the droplets. Indeed, setting *ϵ*_0_ = 0.2 as in our experiments, with typically *L*_0_ = 20 *μ*m and *γ* = 1 mN/m, one finds G’ = 1000 Pa as typically measured with the microplates setup.

The viscous contribution to the complex modulus can be estimated from the drag force F_*v*_ ~ 3*πηVL*_0_. Using the Hertz-like estimation of the droplet-microplate contact area, the stress is expressed as:

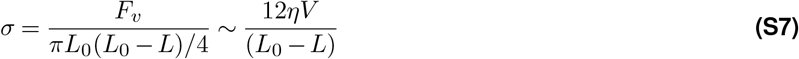

Expressing the typical speed V as function of the rate of strain V = *L*_0_(*dϵ/dt*), one gets:

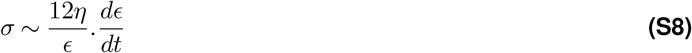

Using the equation **S4**, one finds:

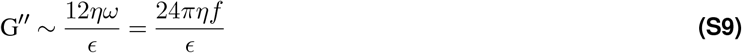

For *f* = 1 Hz, *ϵ*_0_ = 0.2 and the lipiodol viscosity at 37°C *η* = 0.025 Pa.s, one finds G″ (1 Hz) = 10 Pa which is the right order of magnitude (**Fig. 2G**).

As a consequence, comparing *G’* = 1000 Pa and *G″* = 10 Pa, we consider that the viscous contribution can be neglected for processes taking place over time scales longer than a second, and we only take into account the storage modulus over the loss one: *G*\*(*f*) ~ *G*’. Since *G*’ is constant over the explored frequency range (**Fig. 2G**), *G*\*(*f*) ~ *G*’ is thus considered as an apparent Young modulus.

### Supplementary Note 4: Characterization of B cell mechanics

B cell viscoelastic properties was assessed by microplate microrheology. Experiment on single cells show that B cell mechanical properties follow a damping model (**Fig. 2F**). G’ and G″ mainly behave as weak power laws of the frequency (68, 69) (**Fig. 2F**) with a power law exponent similar to those already reported for immune cells (*α* ≈ 0.16), and exhibit an apparent visco-elastic modulus of about 165 Pa (**Fig. 2G**).

### Supplementary Note 5: Characterization of antigen mobility on droplets

An interesting difference between two types of functionalization (bulk vs. surface) is that the B cell is able to gather the antigen only in the first case, despite polarizing correctly in both cases. One possible cause could be a difference in the antigen mobility. We quantified the mobility of the antigen in the two types of droplet by measuring the diffusion coefficient by Fluorescence Recovery After Photobleaching (FRAP) following the protocol described in (61). Diffusion coefficients of bulk- and surface-functionalized droplets are respectively equal to 0.13 ± 0.09 μm^2^/s (N=12) and 0.05 ± 0.03 μm^2^/s (N=12): the two values are significantly different (**Fig. S7**).

Of note, cells known to exert stronger forces, such as macrophages (70), are able to form a cluster of antigens on both types of droplets (24) meaning independently of functionalization process, confirming that the antigen cluster is driven by cell forces. Further molecular dynamic simulations or FRAP experiments out of equilibrium are required to elucidate the physical mechanisms linking pulling forces and active antigen accumulation.

### Supplementary Figures

**Fig. S1.**
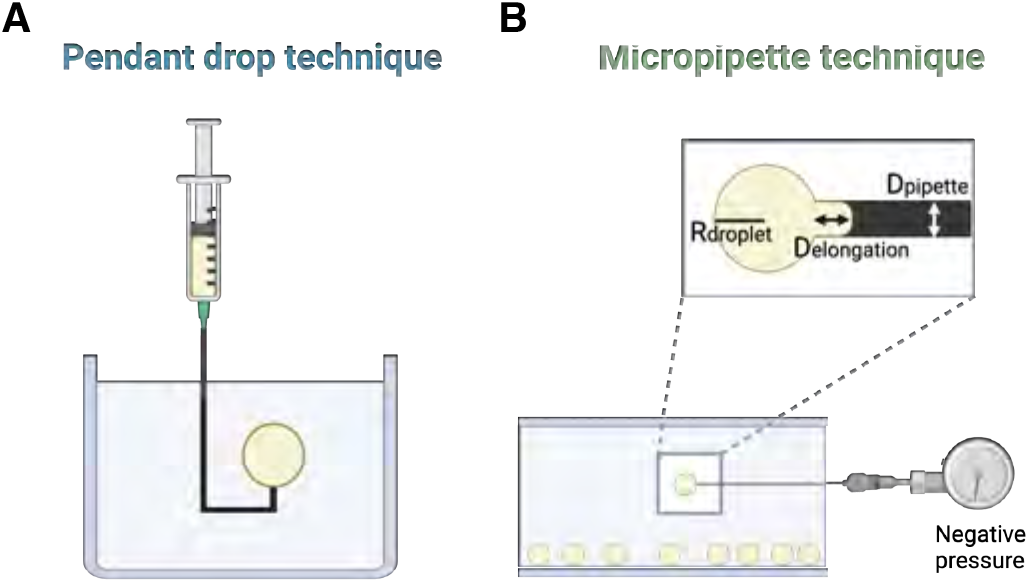
Schematics of pendant drop and micropipette aspiration techniques. (A) The pendant drop technique consists in measuring the interfacial tension of an oil drop in an aqueous phase (water and surfactants). A dedicated software tracks the drop shape and, depending on oil and aqueous phase properties (type and concentration of surfactants and viscosity), computes the interfacial tension over time. (B) Interfacial tension of stabilized droplets is determined by applying a negative pressure on droplets via a micropipette. The interfacial tension *γ* is computed from the negative pressure required to aspire a droplet until forming an elongation whose dimension is equal to the micropipette size, as *γ* = ΔP*R_p_*/2.

**Fig. S2.**
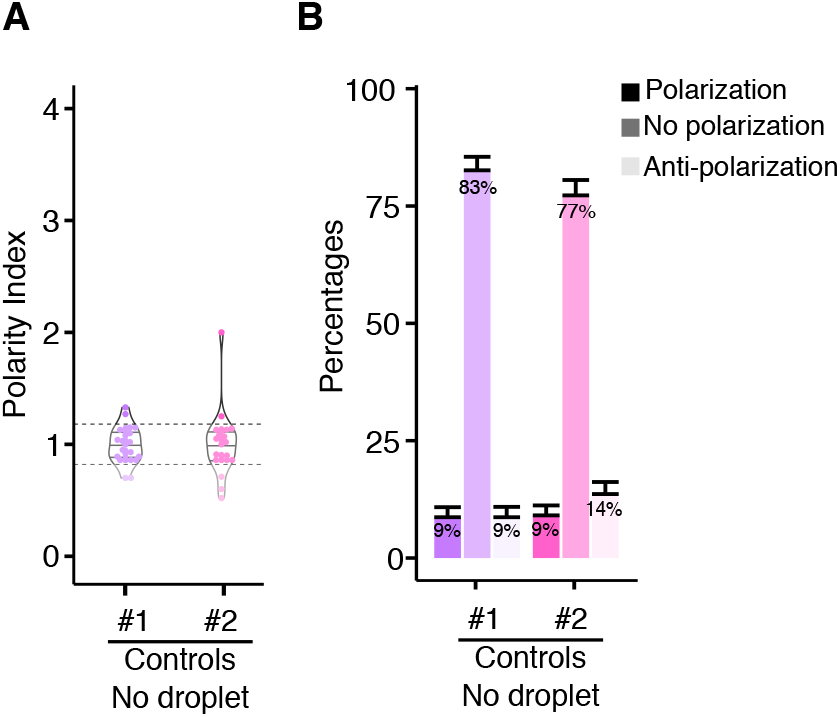
Control of polarization without droplets. Polarity index (left) and percentages of polarized, non polarized and anti-polarized cells (right) without being in contact with droplets. Control #1 has been realized without droplets in the whole chip (N=23 cells, n=3 independent experiments) while control #2 has been realized on non-interacting cells located nearby trapped droplets (N=22, n=3). In both cases, polarization percentage is similar to BSA-coated droplets results.

**Fig. S3.**
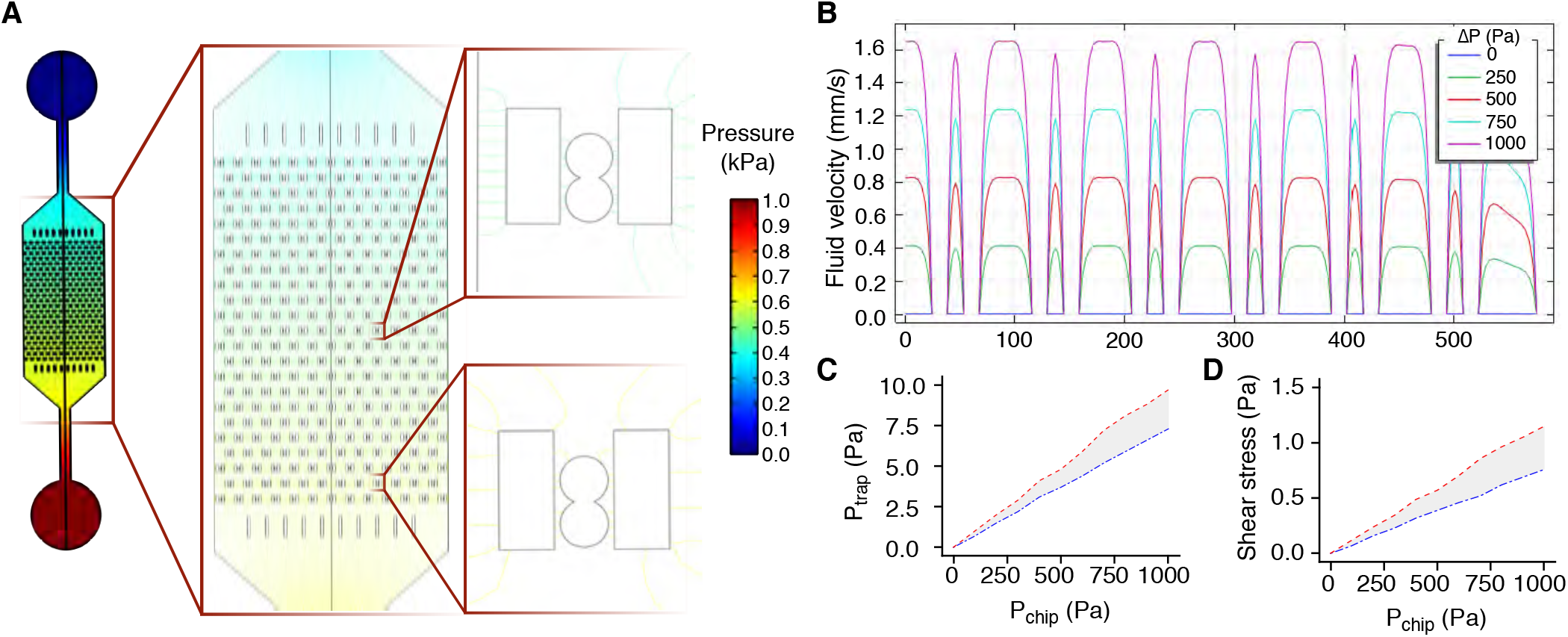
Microfluidic Chip Computational Fluid Dynamics Characterization. (A) Pressure field in the microfluidic chamber for a pressure drop between inlet and outlet about 1 kPa. Each isobar line represents a 1 Pa-step. Depending on trap locations, the pressure variation before and after the trap is between 7.3 and 9.7 Pa. (B) Fluid velocity profiles on a middle cut line depending on the pressure applied at the chip entrance without considering trapped objects, ranging between 0 and 1000 Pa. (C) Pressure drop in a trap depending on the pressure applied in the chip. (D) Shear stress on a trapped cell assumed as rigid, depending on the pressure applied in the chip. (C)-(D) The red curves are obtained for the most mechanically loaded traps, and the blue curves for the least mechanically loaded ones. Simulations were performed with Comsol.

**Fig. S4.**
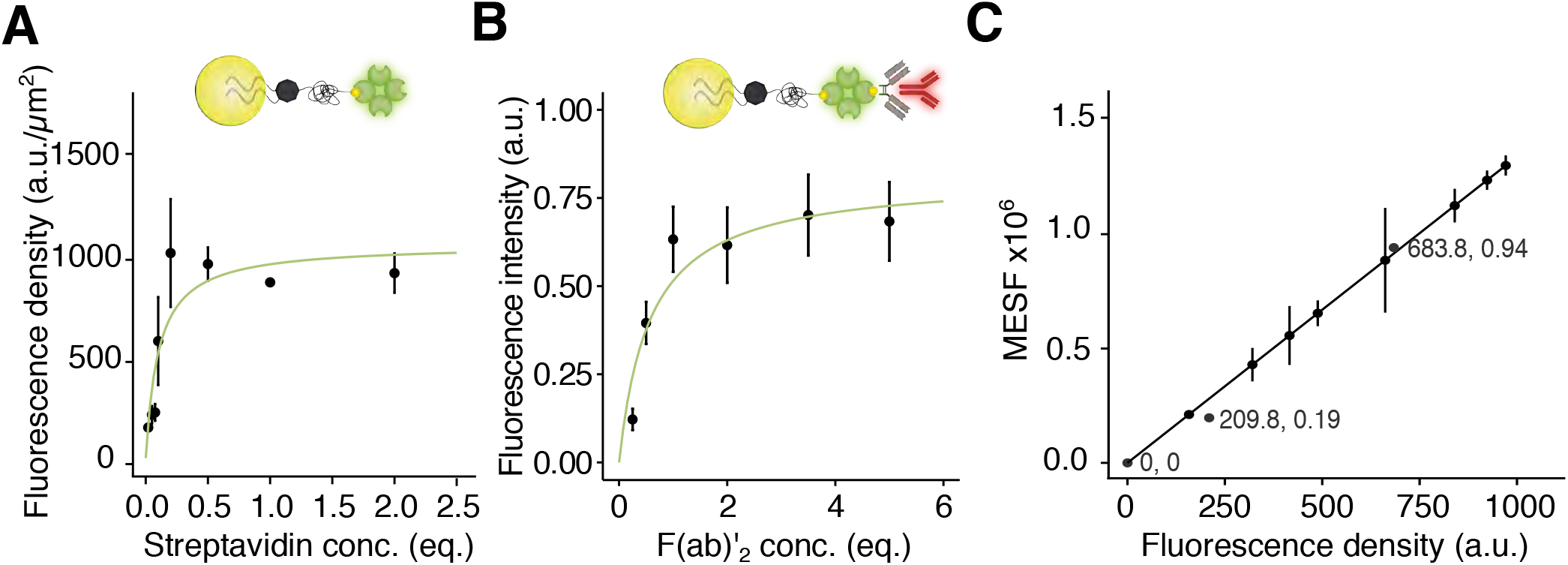
Titration of antigen concentration on droplet surface. (A) Experimental titration curve of streptavidin molecules on 11 *μ*m-large droplets, initially coated with 100 equivalents of DSPE-PEG_2000_-biotin. (B) Experimental titration curve of F(ab’)_2_ fragments on 11 *μ*m large droplets, initially functionalized with 100 equivalents of DSPE-PEG_2000_-biotin and 1 equivalent of Streptavidin. Non-fluorescent F(ab’)_2_ fragments are imaged with a secondary fluorescent anti-F(ab’)_2_ antibody, with a 1:1 ratio. (C) Calibration curve of commercial FITC fluorescent Quantum™ MESF (Bang Laboratories, Inc.) and streptavidin-coated droplets. Fluorescence intensity of beads and droplets have been measured by microscopy (channel GFP). Fluorescence intensity of Quantum™ MESF (dark gray dots) are correlated to MESF provided by the company (coordinates are indicated). MESF of droplets have been determined thanks to the linear function y = 1374.67x.

**Fig. S5.**
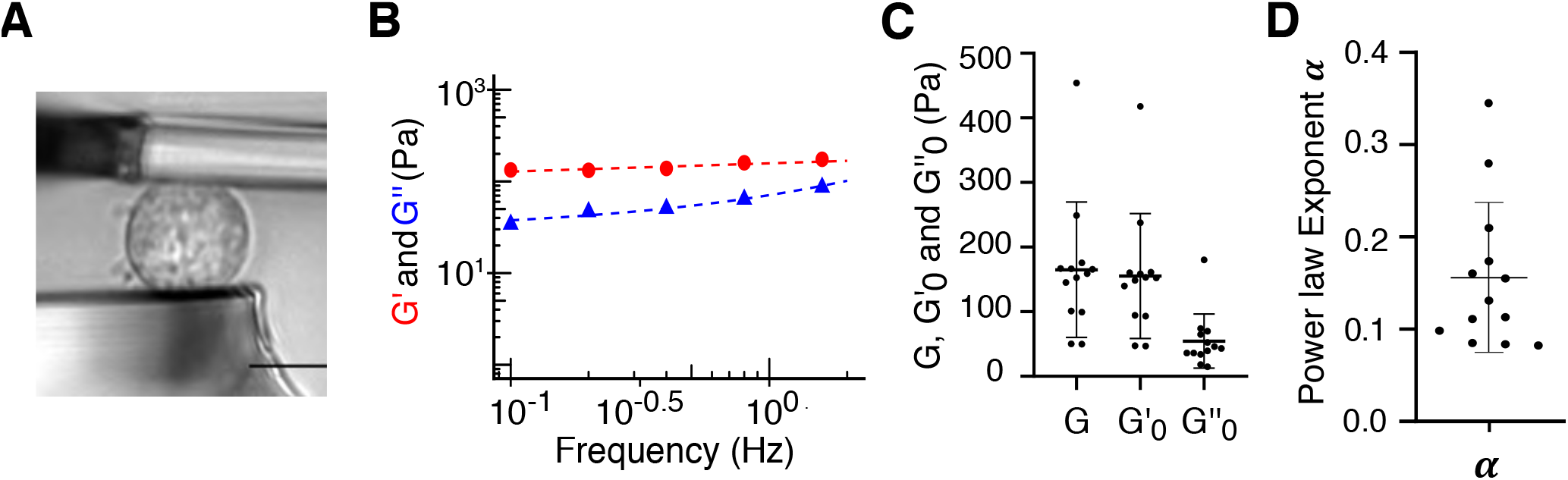
Characterization of B cell mechanics. (A) Representative brightfield image of a B cell immobilized between the microplates. Scale bar: 10 *μ*m. (B) Elastic *G*’ and viscous G″ modulus of a single B cell plotted as a function of the probing frequency. Both G’ and G″ follow a power law. (C) Summary graph of the G’, G″ and G* values of the B cells probed by microplates (N=13 cells, n=3 independent experiments). The average values of 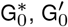 and 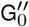 are respectively equal to 165.0± 57.0 Pa, 155.4 ± 52.54 Pa, and 54.5 ± 22.7 Pa. (D) For B cells, the complex modulus G follows a power law of the frequency with coefficient 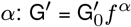 and 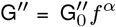. The exponent is determined from fitting of both moduli: *α* = 0.155 ± 0.04 (N=13, n=3 independent experiments), corresponding to values measured for immune cells.

**Fig. S6.**
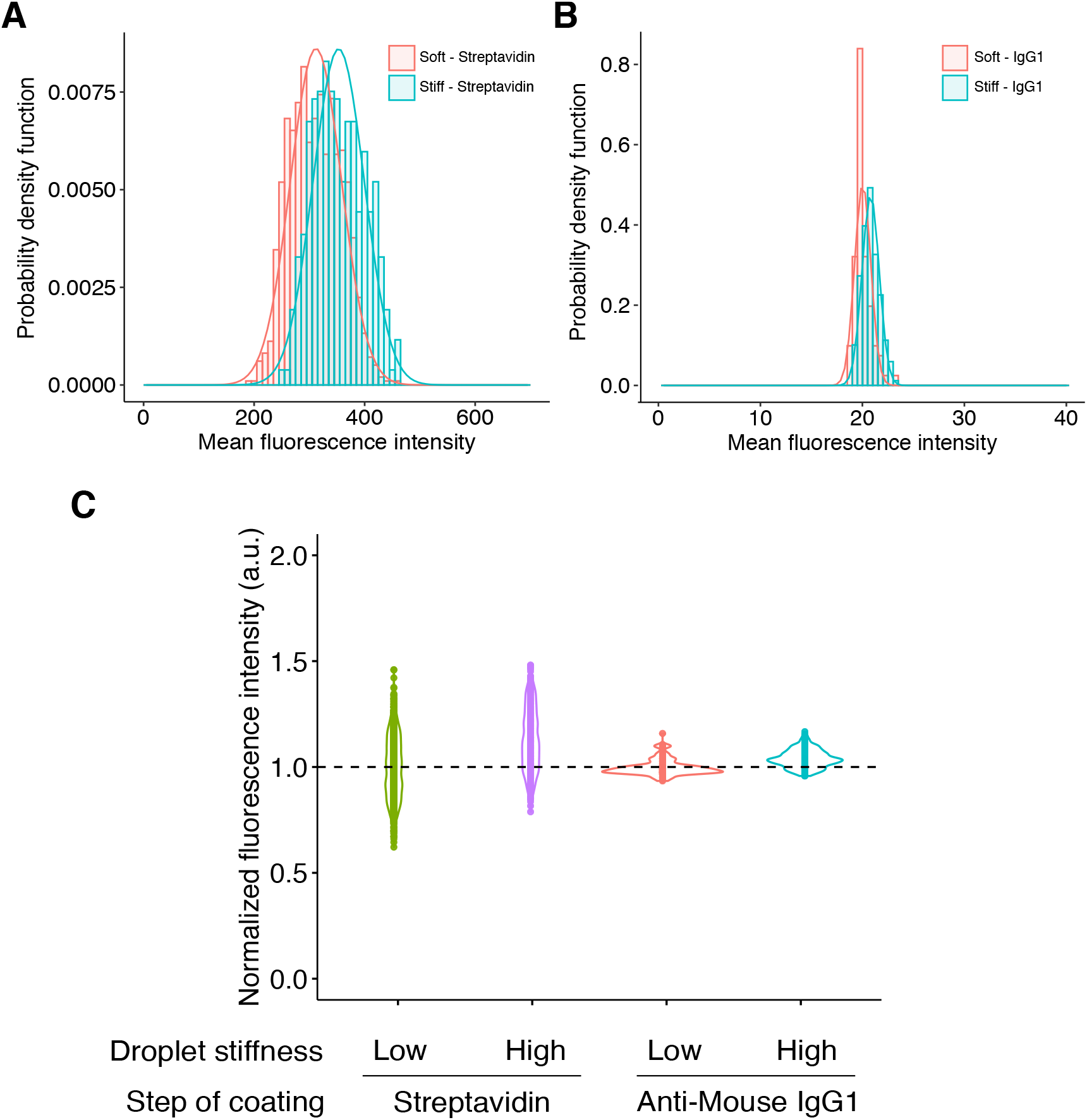
Antigen coating depending on droplet stiffness. Fluorescence intensities of 0.5 equivalent Anti-mouse IgG1 κ Isotype (Alexa Fluor 647) (A) on 1 equivalent of streptavidin (Alexa Fluor 488) (B) bored by soft and stiff droplets. Each histogram is fitted with a gaussian curve with mean ± sd (% sd/mean): values are 19.79 ± 0.79 (4%) (stiffness^*low*^, IgG1), 20.54 ± 0.84 (4%) (stiffness^*high*^, IgG1), 312 ± 46 (15%) (stiffness^*low*^, streptavidin), 353 ± 46 (14%) (stiffness^*high*^, streptavidin). (C) Pooled graph of normalized fluorescence intensities. Intensities relative to stiff droplets are normalized by the mean value of soft droplets for both conditions, IgG1 and Streptavidin.

**Fig. S7.**
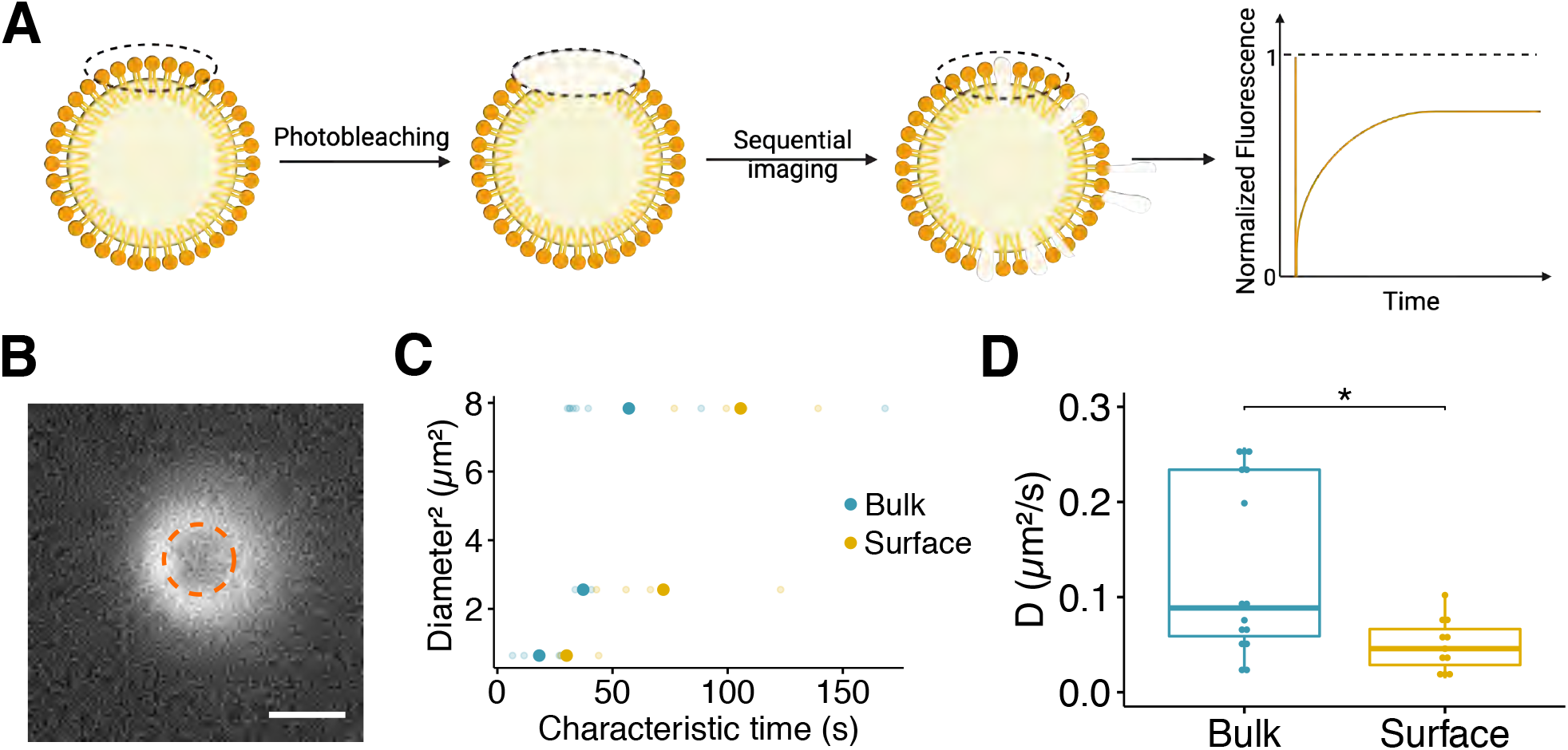
FRAP Experiments - Antigen diffusion. (A) Scheme of the experimental setup. A circular area on to of a single antigen-coated droplet is bleached. The fluorescence recovery is recorded over 5 min and finally plotted to deduce the diffusion coefficient. (B) Confocal image of a bleached functionalized droplet. The bleached area is surrounded by a dashed circle. Scale bar: 5 *μ*m. (C) Correlation of the bleached area and the relative characteristic time for bulk- and surface-functionalized droplets. The shaded dots represent the individual values and the main solid points the mean value. (D) Diffusion coefficients of bulk- and surface-functionalized droplets determined via FRAP experiments, are respectively equal to 0.13 ± 0.09 μm^2^/s (N=15 droplets) and 0.05 ± 0.03 μm^2^/s (N=11 droplets). * indicates p-value<0.05, Wilcoxon test.

**Fig. S8.**
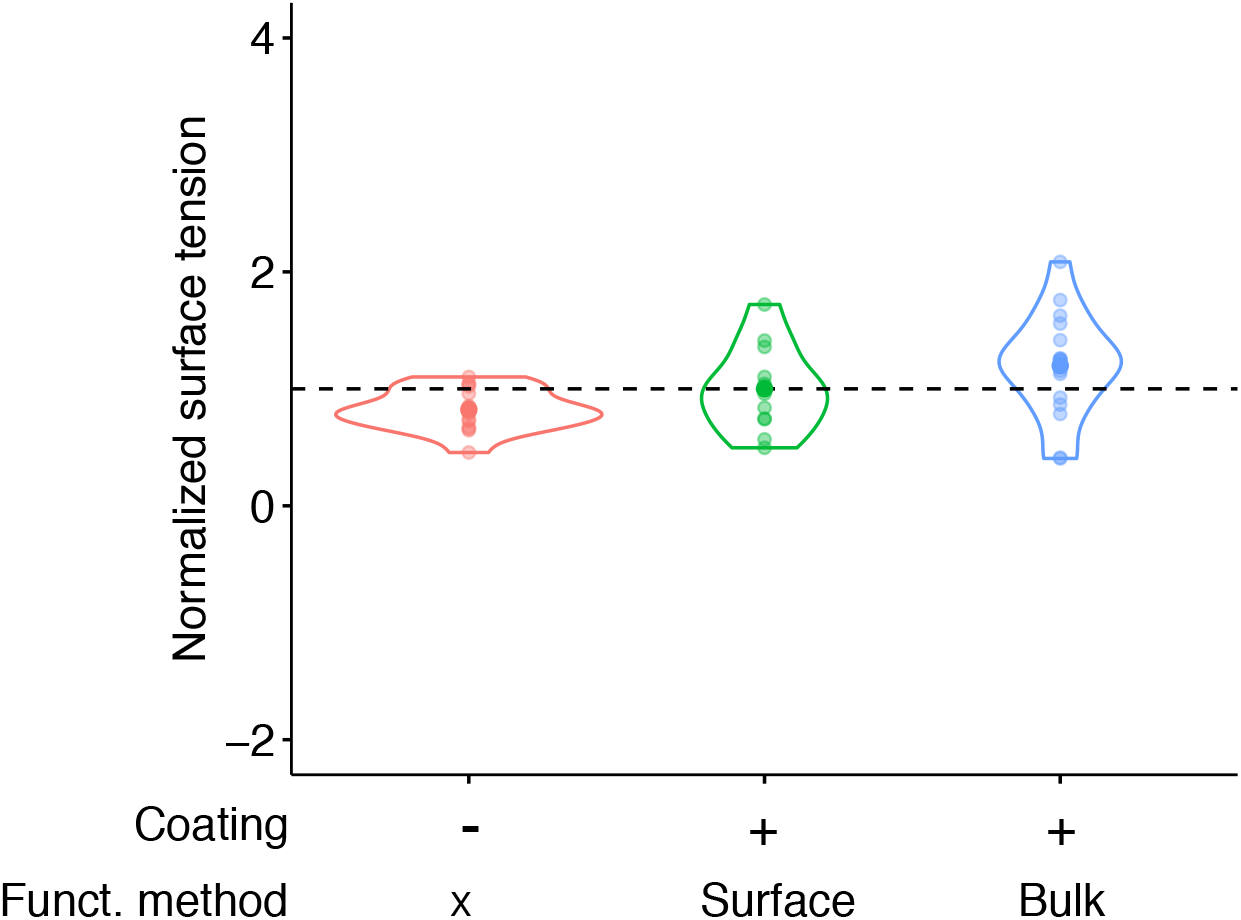
Surface tensions dependance upon functionalization method. Surface tension micropipette measurements of non-coated soybean oil droplets, surface-functionalized and bulk-functionalized F(ab’)_2_-coated soybean oil droplets. Values are normalized by the mean surface tension obtained for the surface-functionalized droplets for sake of clarity. Differences in surface tensions between different coatings and functionalization methods are not significant.

**Fig. S9.**
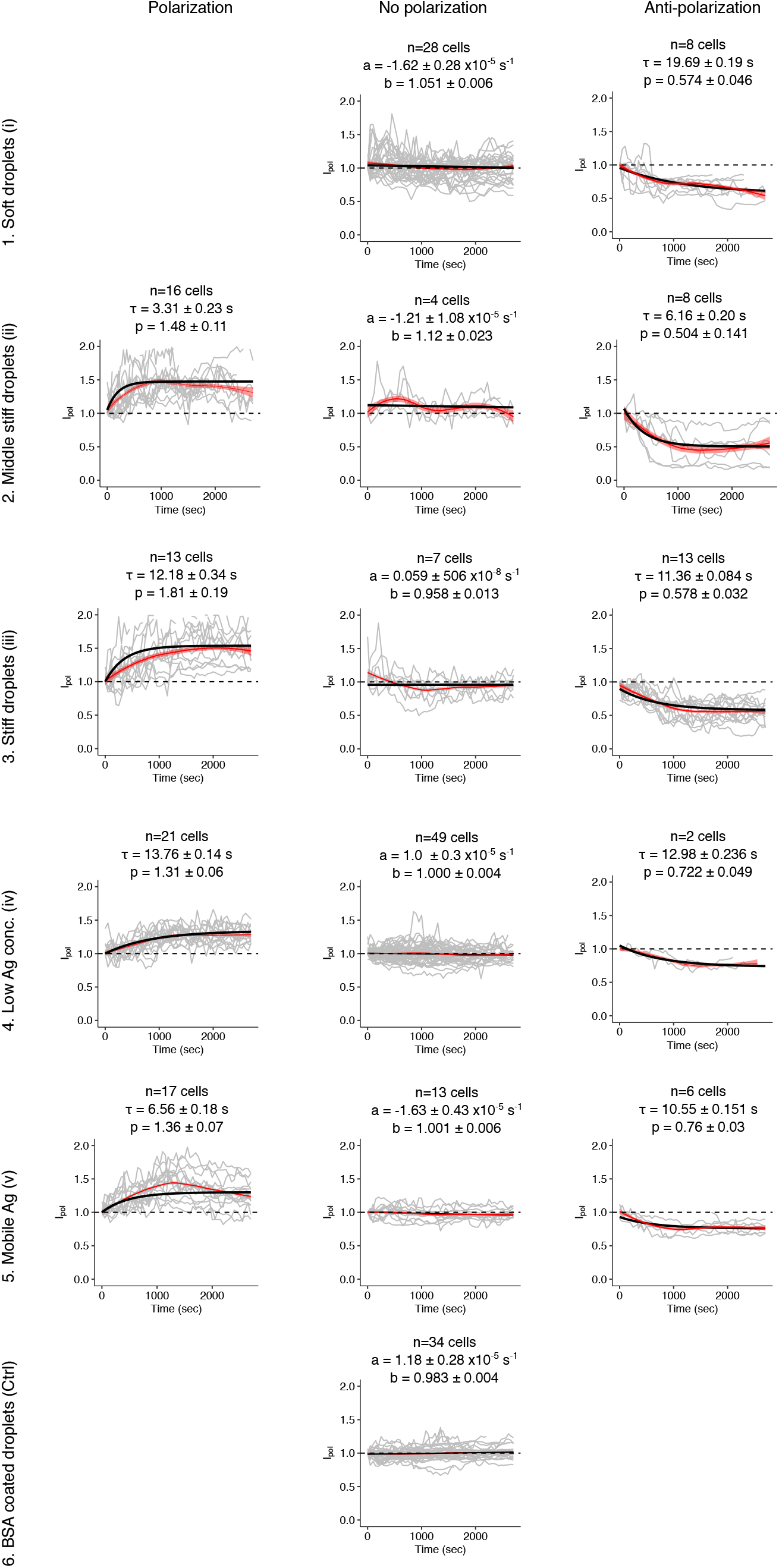
Cellular phenotype of B lymphocytes dynamics. The exponential polarization and anti-polarization fits follow respectively *y_pol_* = (*p* — 1)(1 — exp(-*t/τ*) + 1 and *y_antipol_* = (1 — *p*)(1 — exp(—*t/τ*)) +1, where *τ* is the characteristic time and *p* the related plateau. The non-polarized phenotype is characterized by a linear fit: *y_nopol_* = *ax* + *b*.

**Fig. S10.**
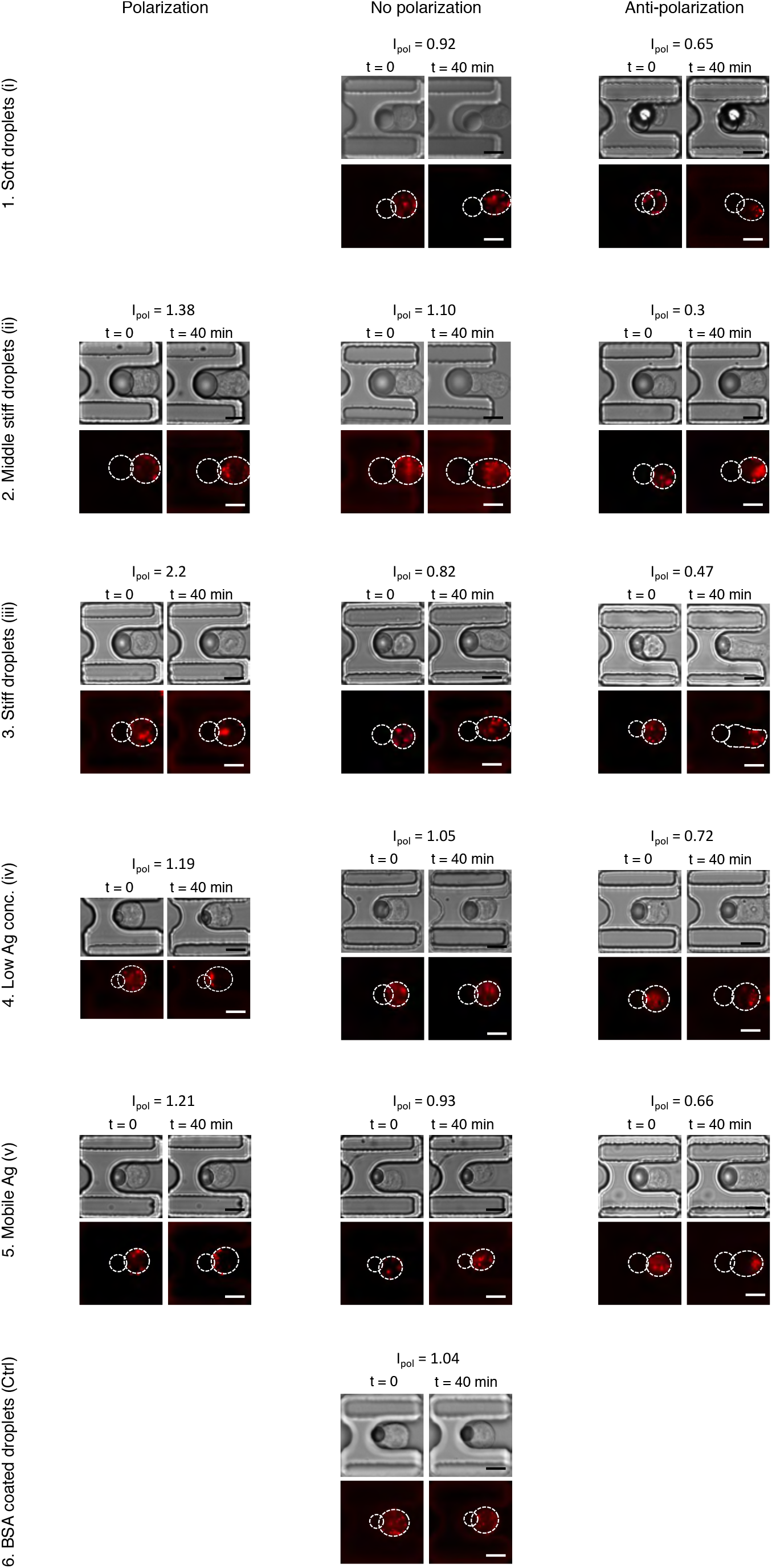
Microscopy image of B lymphocytes dynamics. Images show the first time point (contact) and the final one (40 minutes) for different droplet conditions and for the three behaviors (polarization, no polarization, and anti-polarization). Scale bar: 10 *μ*m.

### Supplementary Tables

**Table S1.** Interfacial tension of oil/aqueous phases via pendant drop technique.

| Oil phase | Water Phase | Surface tension (mN/m) |
| --- | --- | --- |
| Mineral oil + Oleic acid 5%v/v | F68 15% | 1.69 ± 0.19 |
| Mineral oil | F68 15% | 4.73 ± 0.38 |
| Soybean oil | F68 15% | 12.03 ± 0.52 |
| Lipiodol oil | F68 15% | 1.52 ± 0.12 |
| Lipiodol oil + Oleic acid 5%v/v | F68 15% | 0.88 ± 0.12 |

**Table S2.** Interfacial tension of oil/aqueous phases via micropipette technique. Measurements have been performed on droplets stabilized by Pluronic F68 and surrounded by an aqueous phase (Phosphaste buffer/Tween 20 at CMC). *From (71)

| Oil phase | Functionalization | Surface tension ± SD (mN/m) |
| --- | --- | --- |
| Lipiodol oil + Oleic acid 5%v/v | None | $2.36 \pm 0.31$ |
| Lipiodol oil + Oleic acid 5%v/v | DSPE-PEG-Biotin | $2.50 \pm 0.24$ |
| Soybean oil | DSPE-PEG-Biotin | $10 \pm 2^*$ |

**Table S3.** Table depicting the equivalent and the volume required to coat 2 millions of 12 *μ*m-large droplets, via surface functionalization method.

| Functionalization | Equivalent | Conc.(mg/ml) | Volume (μL) |
| --- | --- | --- | --- |
| DSPE-PEG(2000)-Biotin | 100 | 10 | 3.60 |
| Streptavidin | 1 | 1 | 5.43 |
| Biotinylated F(ab')<sub>2</sub> | 1 | 1.4 | 0.98 |
| Secondary Antibody | 1 | 0.8 | 2.35 |

## Supplementary Movies

Movie S1: Active recruitment of antigen at the B cell (IIA1.6) synapse in contact with a F(ab’)_2_-coated bulk-functionalized stiff droplet. 40X objective, epifluorescence microscope, LysoTracker DND staining. Timeframe: 50 s.

Movie S2: Polarized B cell IIA1.6 in contact with a F(ab’)_2_-coated surface-functionalized stiff droplet. 40X objective, epifluorescence microscope, LysoTracker DND staining. Timeframe: 50 s.

Movie S3: Anti-Polarized B cell IIA1.6 in contact with a F(ab’)_2_-coated surface-functionalized stiff droplet. 40X objective, epifluorescence microscope, LysoTracker DND staining. Timeframe: 50 s.

## Footnotes

1 This number comes from an estimation that considers: 20000 receptors (IgM in primary cells) (51) squeezed in a synaptic region of radius 7*μ*m: 20000/*π*7^2^ ≈ 130 mol/*μ*m^2^ interacting with bivalent ligands.

